# Automated highly multiplexed super-resolution imaging of protein nano-architecture in cells and tissues

**DOI:** 10.1101/791053

**Authors:** Maja Klevanski, Frank Herrmannsdoerfer, Varun Venkataramani, Steffen Sass, Mike Heilemann, Thomas Kuner

**Affiliations:** Department of Functional Neuroanatomy, Institute for Anatomy and Cell Biology, Heidelberg University, Im Neuenheimer Feld 307, 69120 Heidelberg; Institute of Physical and Theoretical Chemistry, Goethe-University Frankfurt, Max-von-Laue-Str. 7, 60438 Frankfurt/M

## Abstract

Understanding the nano-architecture of protein machines in diverse sub-cellular compartments remains a challenge despite rapid progress in super-resolution microscopy. While singlemolecule localization microscopy techniques allow the visualization and identification of cellular structures with near-molecular resolution, multiplex-labeling of tens of target proteins within the same sample has not yet been achieved routinely. However, single sample multiplexing is essential to detect patterns that threaten to get lost in multi-sample averaging. Here, we report maS^3^TORM (**m**ultiplexed **a**utomated **s**erial **s**taining **s**tochastic **o**ptical **r**econstruction **m**icroscopy), a microscopy approach capable of fully automated 3D dSTORM imaging and solution exchange employing a re-staining protocol to achieve highly multiplexed protein localization within individual biological samples. We demonstrate 3D super-resolution images of 15 target proteins in single cultured cells and 16 targets in individual neuronal tissue samples with <10 nm localization precision. This allowed us to define novel nano-architectural features of protein distribution within the presynaptic nerve terminal.

## INTRODUCTION

Proteins in cells interact as networks in space and time. Knowing the molecular organization of proteins relative to each other is crucial to understand mechanisms of cellular function. This advance requires (i) imaging techniques that are capable to visualize cellular structures with near-molecular resolution, (ii) strategies for multi-protein staining of an individual biological sample, and (iii) an automated workflow for non-invasive solution exchange and data acquisition. Identifying complex nano-architectural patterns of 3D protein arrangement requires the identification of as many as possible different proteins within a single sample, because averaging distribution patterns determined from many individual samples may mask the underlying organizational principle, at least for structures that are not inherently symmetrical.

Optical super-resolution microscopy has revolutionized cell biology (reviewed in Sigal et al. 2018^1^). One branch of techniques is single-molecule localization microscopy (SMLM)^2^, comprising (direct) stochastic optical reconstruction microscopy ((*d*)STORM)^3,4^, photoactivated localization microscopy (PALM)^5^, points accumulation for imaging in nanoscale topography (PAINT)^6^, and DNA-PAINT^7^. SMLM achieves a localization precision of below 10 nm, enabling to decipher protein nano-architecture in 3D, and intrinsically providing access to molecule numbers^8^.

Multiplexed protein labeling remains a challenge to all fluorescence microscopy methods. Stationary labels are restricted owing to spectral limitations, on the one side, and lack of efficient elution protocols, on the other. Only few multi-color SMLM reports exist so far^9–11^. Dynamic labels, such as reversibly binding fluorophore-labeled DNA strands as used in DNA-PAINT, are ideally suited for multiplexed imaging^12,13^. However, multiplexing with DNA-PAINT is restricted to available primary labels, such as antibodies or aptamers^14^. Multiplex imaging at large scale inevitably requires automated solution exchange, sample positioning, image acquisition, and re-staining techniques. For automation, Almada et al. (2018)^15^ employed the NanoJ fluidics system to perform a five-color experiment. Another recent approach successfully demonstrated high-throughput STORM screening of cells in multiwell plates^16^. However, this approach does not allow multiplex analysis within individual samples. To date, a fully automated microscopy system including non-invasive solution exchange is not available.

The nano-architecture of active zones (AZs), presynaptic specializations mediating neurotransmitter release, has been addressed in studies mostly focusing on the molecular architecture in the *Drosophila* neuromuscular junction^17^. Mammalian central synapses and their nano-architecture have been studied in synaptosomes^18^ and cultured hippocampal neurons^19–22^. However, only few studies addressed presynaptic protein topography in mature synapses of nervous tissue, such as the calyx of Held ^23^, using super-resolution microscopy ^24–26^. A systematic investigation of molecular nano-architecture in natively matured synapses has not yet been achieved. SMLM imaging of neuronal tissue^25,27–29^ and even more multiplexing remain challenging because of technical limitations.

Here, we developed a universal method for serial detection of multiple targets that is suitable for cells and tissues. Our fully-automated system comprises an in-house built 3D *d*STORM setup, a pipetting robot, and the custom-written software allowing coordinated operation of both systems. We illustrate that maS^3^TORM can execute re-staining protocols to perform multiplex experiments with at least 15 targets that could be detected in single cells and 16 targets in neuronal tissue, allowing us to study the protein landscape within individual synapses. For example, the localization of motor protein myosin Va (MyoVa) is correlated with synaptic vesicle (SV) marker VGlut 1 throughout the synapse, suggesting that in addition to its role in SV trafficking, MyoVa could play a role in SV clustering and tethering. Furthermore, we observed that polymerized actin is less abundant in close proximity to membranes with AZs compared to AZ-free membrane stretches, in line with observations that actin dynamics regulates neuronal activity^22,30–32^.

## RESULTS

### maS^3^TORM setup and multiplexing workflow

maS^3^TORM is composed of a home-built *d*STORM setup and the commercially available pipetting robot PAL3 RTC (**Fig. 1a, b; Supplementary Fig. 1** and **Supplementary Video 1**). The setup is designed for astigmatism-based 3D measurements^33^ and allows simultaneous acquisition of two channels by spectral demixing (**Supplementary Fig. 1a**)^34^. For axial drift correction, it is equipped with a focus stabilization module (**Fig. 1a, b, Supplementary Fig. 1a**). To ensure an automated and coordinated interplay between the staining procedures carried out by the pipetting robot and image acquisition, the Experiment Editor software was developed that can address both the μManager^35^ based Microscope Control software and the commercial PAL3 Chronos software (**Supplementary Fig. 1b**). In the Experiment Editor, the user can create customized experimental workflows by combining microscope- and robot-related modules and iterating tasks for a desired number of times (**Supplementary Fig. 2**).

**Figure 1.**
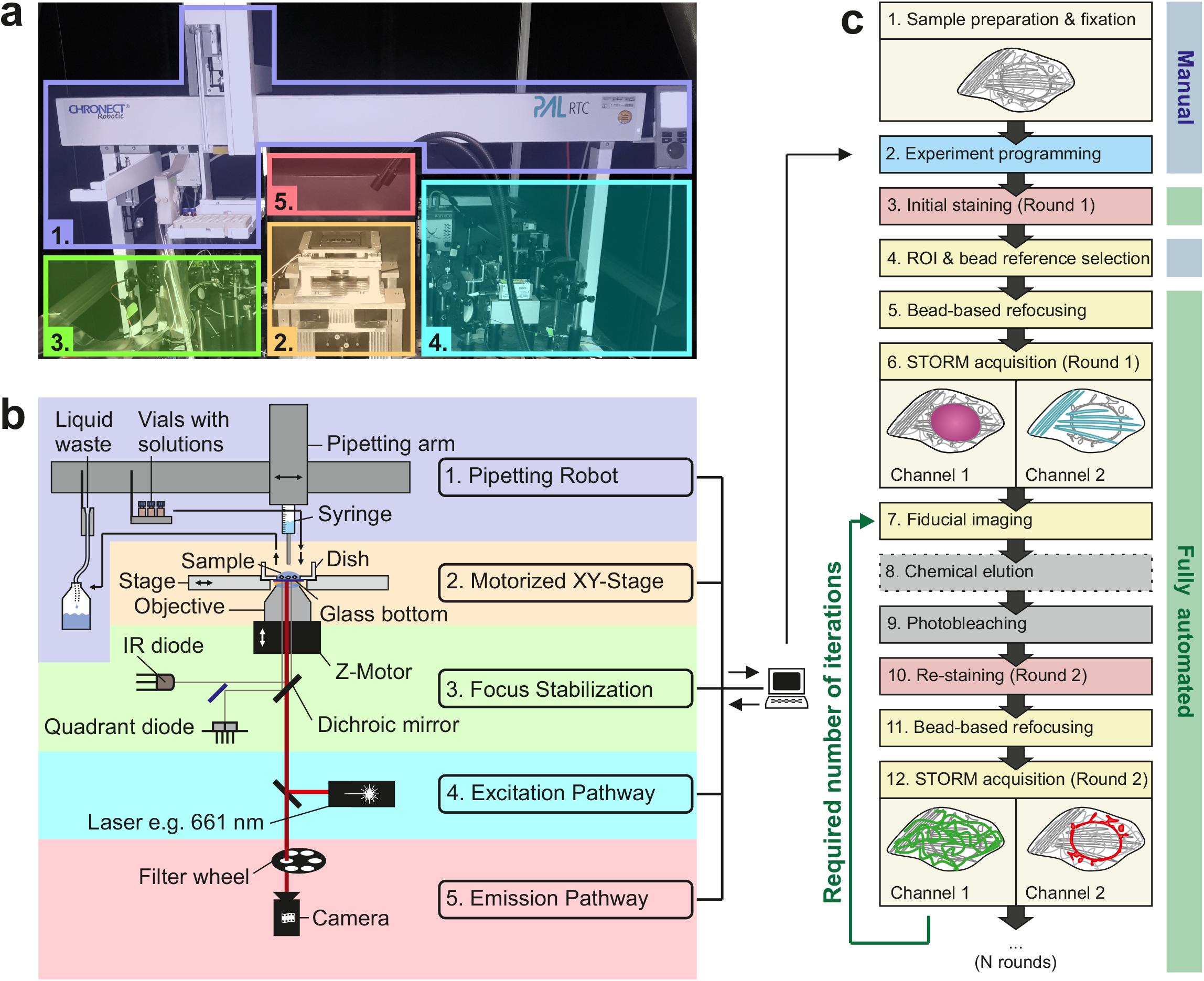
Automated maS^3^TORM multiplex setup and experimental workflow. Photographic (**a**) and schematic overview (**b**) of the setup body comprising the pipetting robot (1, purple), the motorized stage on top of the inverted 1.49 NA 100× objective (2, yellow), the focus stabilization system (3, green), components of the excitation (4, blue), and the emission pathway (5, red). (**c**) Experimental workflow for an automated multiplex approach. From step 5 onwards the workflow is fully automated. Step 8 is conditional (stippled line). Details on the series of steps in an automated imaging session see Methods.

For multiplex experiments, we developed re-staining workflows allowing fixed cells or tissues to be serially labeled, imaged and stripped (**Fig. 1c**). Once the exact experimental workflow is designed and programmed using the Experiment Editor, the designated vials of the PAL3 pipetting robot can be filled with respective solutions and the fixed sample can be placed onto the motorized stage. After the initial staining is performed via Experiment Editor, regions of interest (ROIs) are selected manually and their coordinates are entered into the predesigned workflow in Experiment Editor using a specialized ROI module. To make sure that the initially selected focal plane can be retrieved in each imaging round, we developed an auto-refocusing system (for details see Methods) that is based on imaging of a reference region with fluorescent TetraSpeck beads. Next, dual-color 3D *d*STORM data of the preselected ROIs are acquired and automatically redirected and processed by rapidSTORM^36^. For precise registration of images from distinct staining rounds, we imaged fiducial beads that were seeded underneath cells or tissue for maximal immobilization. We found a way to preserve fluorescence of the fiducials for at least 11 imaging and bleaching rounds by moderately exciting them using the 561 nm laser while recording through the 700/75 bandpass filter (same filter as used for *d*STORM acquisition) for minimal chromatic aberration. Feature-based lateral drift correction within individual superresolution images was realized in subsequent post-processing analysis.

Depending on the type of labels used, signal removal was achieved by photo-bleaching using combined exposure to 405 nm and 661 nm laser^10^ or a combination of chemical elution with 0.1% SDS at pH 13 (modified from Collman et al. (2015)^37^) and photo-bleaching (for details see Methods). Control experiments with U2OS cells labeled with Tom20 antibody showed that bleaching successfully removed 98.9 ± 0.9% of the initial signal (**Supplementary Fig. 3**, Supplemental Note 1). However, if primary antibodies of the same species are used in consecutive staining rounds bleaching is not sufficient. In this case antibodies need to be removed physically by elution in addition to bleaching the fluorophores. After elution and bleaching, only 1.8 ± 1% of the initial Tom20 signal can be retrieved by re-application of the secondary antibody in the presence of a competing primary antibody (**Supplementary Fig. 3**). This shows that our strategies for signal removal cause minimal crosstalk between staining rounds (for examples with other targets, see **Supplementary Fig. 4a-d**), a prerequisite for high-fidelity multiplexed imaging. Moreover, staining can be maintained while many additional staining rounds are added (**Supplementary Fig. 4e**), another essential condition to maximize the number of localizations in the same sample.

### Multiplexed 3D super-resolution imaging of a single eukaryotic cell

We next demonstrate multiplexed 3D *d*STORM imaging in fixed U2OS cells. Within 56 h, we performed 11 rounds of staining, and extracted 15 high-quality super-resolution data sets for further analysis (**Fig. 2a**; see **Supplementary Table 1** for a detailed workflow). The localization precision of SMLM images shown in **Figure 2a** remained stable at a high grade with a mean value of 7.7 ± 2.7 nm (calculated according to the nearest neighbor distance method^38^; **Supplementary Fig. 5**). The targets comprised Nup133 of the nuclear pore complex, F-actin, sialic acid and N-acetylglucosaminyl residues (bound by WGA) amongst others present at the cell membrane and in the center of nuclear pores, CHC17 of clathrin-coated vesicles, α-tubulin, paxillin in focal adhesions, intermediate filament vimentin, GM130 in Golgi, fibrillarin in nucleoli, Tom20 in mitochondria, lamin A/C at the nucleus, PDI in the ER, Rab5 and EEA1 in early endosomes, and DNA labeled by JF_646_-Hoechst conjugate^39^. All images could successfully be registered to each other with good precision (**Fig. 2b**). The quality of the images and the registration process are illustrated in **Figures 2c, d** showing Nup133 signal localized at the peripheral subunits of nuclear pores merged with WGA signal in the center of nuclear pores. The 3D views reveal that the Nup133 signal originates from two layers/rings of the nuclear pore complex (lower right panel in **Fig. 2d**). All data were collected in 3D mode and can be registered along all three axes as exemplified by α-tubulin and vimentin that partially run parallel to each other but do not colocalize (**Fig. 2e**; for other 3D examples see **Supplementary Fig. 6** and **Supplementary Video 2**). Finally, magnified *d*STORM images of endosome-related and - adjacent structures (**Fig. 2f**) prove that early endosomes can be successfully re-stained by two different endosomal markers, Rab5 and EEA1. As expected, markers for Golgi, clathrin-coated structures, and associated actin patches^40,41^ can be found in close proximity. All the above examples demonstrate a robust, multi-target super-resolution imaging using maS^3^TORM.

**Figure 2.**
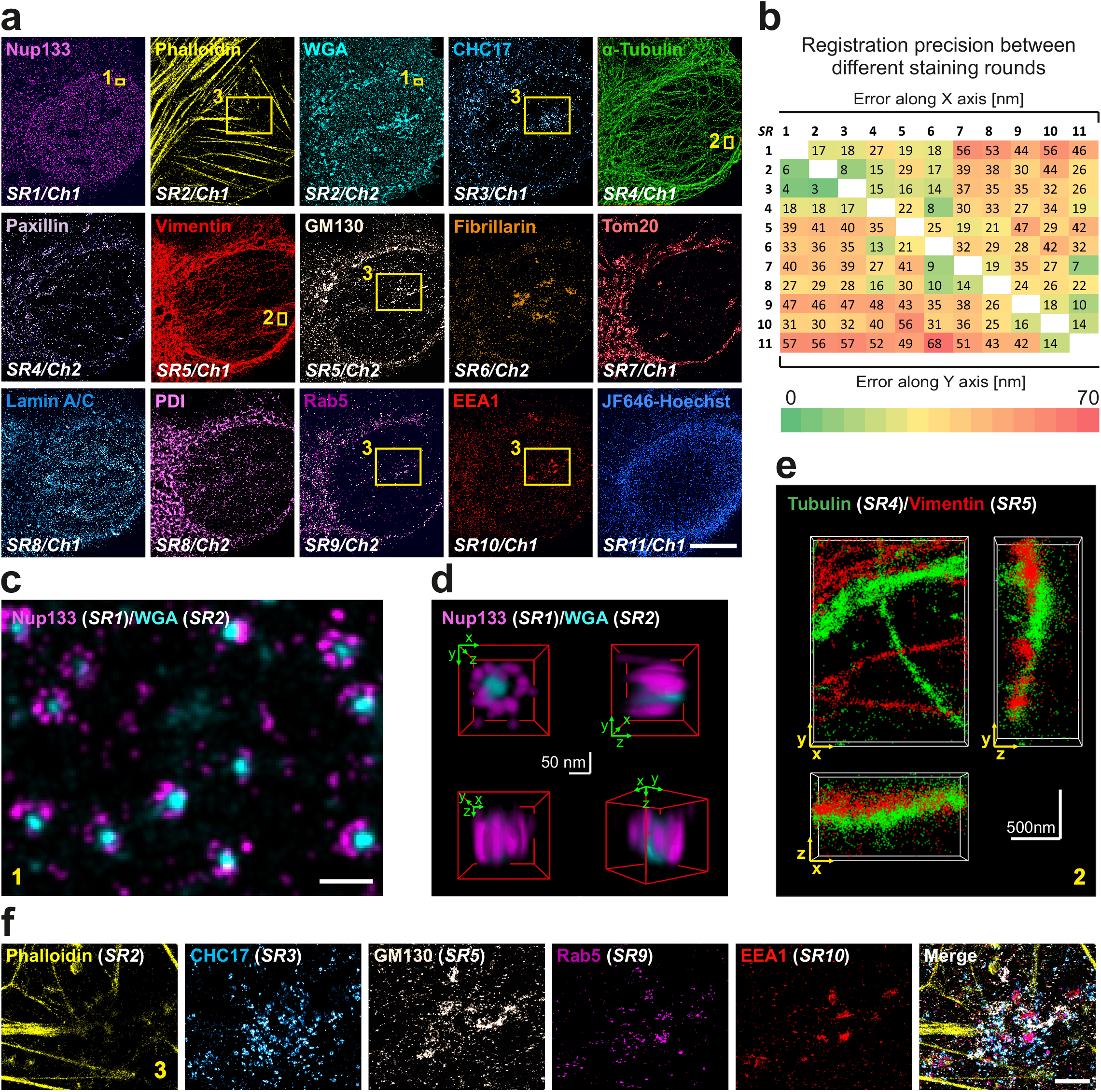
Multi-target 3D STORM of a single U2OS cell. (**a**) A U2OS cell with 15 targets labeled and acquired by 11 staining rounds (SR). Two channels (Ch) can be detected per SR, but not always each channel can be used for final analysis. In addition, to get optimal signal for some sensitive labels (e.g. GFP nanobodies against the Ypet-tag of Nup133), it was beneficial to not stain targets in the second channel. Boxes denote regions magnified in **c** and **f**. Further details see Methods section “Imaging experiments on U2OS cells”. (**b**) Heatmap representing the registration error along the X (upper matrix part) and the Y axis (lower matrix part) reveals precise registration, in particular, between images of adjacent staining rounds (arranged along the white diagonal fields). (**c**) Magnified view of boxed region 1 in a with merged peripheral Nup133 and central WGA nuclear pore labeling demonstrating that super-resolution images from consecutive SRs can be registered with high precision. (**d**) 3D views of a nuclear pore complex (from the multiplex experiment in **a**) demonstrate the high lateral and axial resolution and registration precision. (**e**) Merged tubulin and vimentin 3D images from consecutive SRs demonstrating that super-resolution images can be acquired and registered in 3D (boxed region 2 in a). (**f**) Boxed region 3 in **a**): phalloidin-labeled actin filaments, CHC17-labeled clathrin-coated structures, GM130-labeled proximal Golgi apparatus, endosome-associated proteins Rab5 and EEA1, five color merge. Scale bars correspond to 10 μm in **a**, 200 nm in **c**, 50 nm in **d**, 500 nm in e, and 2 μm in **f**.

### Multiplexed imaging of 16 presynaptic proteins in neuronal tissue

We next applied multiplex imaging to brain tissue containing a well-established glutamatergic model synapse referred to as the calyx of Held^23^. The giant presynaptic calyx of Held envelopes the postsynaptic principal cell (**Fig. 3a**) and forms swellings surrounding the oval-shaped postsynaptic cell in a ring-like arrangement (cross-section in middle panel) with AZs situated at the synaptic calyx border (lower panel). **Figure 3b** illustrates the arrangement of four calyces as shown in **Figure 3c**. Tissue sections with a thickness of 400 nm were obtained using a cryo-ultramicrotome and allowed for direct access of antibodies to the target sites without the use of detergents (see Methods). In 10 consecutive staining rounds we were able to super-resolve the distribution of the following 16 targets *in situ* (**Fig. 3c**): AZ marker bassoon, postsynaptic density marker homer1/2/3, F-actin, sialic acid and N-acetylglucosaminyl residues amongst others present at the cell membrane and in Golgi, CHC17 in clathrin-coated structures, MAP2 at the postsynapse, SV marker VGlut 1, GM130 in Golgi, presynaptic Rab3a, Tom20 in mitochondria, α-tubulin, γ-actin, SV marker synaptophysin 1, α/β-synuclein, PDI in ER, and DNA labeled by JF_646_-Hoechst. Individual SMLM images could be registered with excellent precision (**Fig. 3d**). These data demonstrate the feasibility of imaging the distribution of 16 individual protein targets within a single tissue volume at super-resolution.

**Figure 3.**
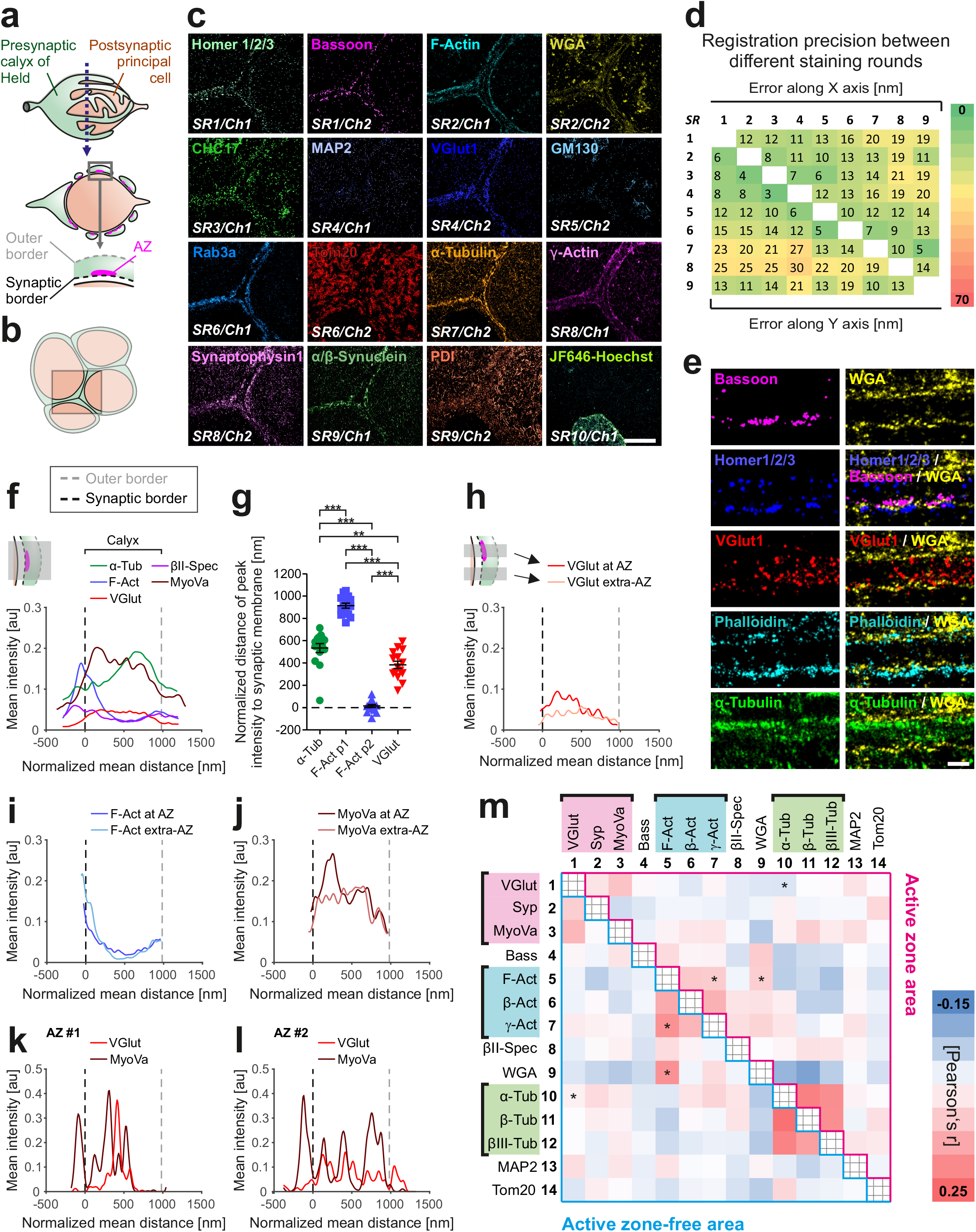
Multiplex STORM of the calyx of Held synapse. (**a**) Schematic representation of the calyx of Held giant synapse (upper panel). The diameter along the stippled line is 15 to 20 μm. Cross-sections through the middle of the synapse (middle panel) revealing multiple swellings each containing multiple active zones (AZ). Zoom-in view (lower panel) of the cross-section showing the outer (gray dashed line) and the inner, postsynapse-proximal, (black dashed line) membrane of the presynapse containing an AZ. (**b**) Cartoon illustrating the topology of adjacent calyces as represented in **c**. (**c**) Re-STORM images of a single brain section containing calyx synapses showing 16 targets acquired in 10 consecutive staining rounds (SR) using one or two spectral channels (Ch). Further details see Methods section “Imaging experiments at the calyx of Held giant synapse”. (**d**) High registration precision (6-30 nm) of images shown in **b** achieved throughout all staining rounds. (**e**) Magnified images of a calyx cross-section (as depicted in **a**, lower panel). Calyx borders labeled with WGA. A sandwich appearance of Homer, Bassoon and WGA staining in the merged image confirms that the membranes of the calyx cross-section are oriented perpendicularly to the imaging plane. (**f**) Presynaptic distribution of selected proteins analyzed and averaged over 9-15 calyces from 3-5 experiments. Synaptic (corresponding to 0 nm at the x-axis) and outer calyx borders (ca. 1000 nm) were determined based on WGA staining (see Methods). (g) The distances of the α-Tub peak, F-Act peak 1 (p1), peak 2 (p2) and VGlut 1 peak to the synaptic membrane differed significantly (**, p ≤ 0.01; ***, p ≤ 0.001, oneway ANOWA). (**h-j**) Differential distribution of selected proteins within calyx regions containing active zones (AZs) and regions lacking AZs (AZ-free) (see inset in **h**). (**h**) Synapse-proximal regions contain more SVs (visualized by VGlut 1 antibodies) at AZ sites as compared to extra-AZ regions. (**i**) Less polymerized actin in AZ-proximal region. (**j**) More MyoVa at AZ sites, correlating with the VGlut 1 profile. (**k, l**) Example of individual AZs supports a correlation between SV and MyoVa distribution and illustrates that individual AZs nevertheless differ from each other. (m) Colocalization matrix showing Pearson’s r coefficients (for exact r values, see **Supplementary Figure 9**) between all combinations of the 14 analyzed proteins at AZ-proximal (upper matrix part) and AZ-free (lower part) calyx regions. Scale bars correspond to 5 μm in **c** and 500 nm in **e**. For sample numbers, see **Supplementary Table 5**.

### maS^3^TORM uncovers global and local protein sociology at the calyx of Held synapse

For quantitative analyses of selected cytoskeletal and synaptic proteins at the calyx of Held, we conducted five independent multiplex experiments. As our main reference for synaptic geometry we used the wheat germ agglutinin (WGA) marker that proved to efficiently label the calyceal borders. Close-up *d*STORM images in **Figure 3e** demonstrate that WGA signal at the membrane facing the postsynapse is sandwiched between presynaptic bassoon and postsynaptic homer. As expected, synaptic vesicle (SV) marker VGlut 1 fills the presynaptic space indicated by WGA. Exemplary images suggest that actin filaments show a rather peripheral localization while microtubules can be found further inside the calyx. These observations are also reflected in the averaged line profiles (**Fig. 3f**) laid across the calyx lumen perpendicularly to the presynaptic membrane. Average plots were obtained from a total of 15 calyces imaged in 5 independent experiments. For statistical reliability, 4-16 line profiles per calyx were analyzed. Calyceal borders were determined based on the WGA signal and all data were normalized according to the mean calyx thickness of 987.2 ± 238.6 nm (with 0 nm corresponding to the inner synaptic and 987.2 nm to the outer border). For details on line profile analysis, see Methods and **Supplementary Figure S7**. Interestingly, βII-spectrin shows a similar spatial distribution as F-actin, and myosin Va (MyoVa) resembles the spatial distribution of VGlut 1. Quantitative analysis of the spatial patterns of α-tubulin, F-actin (two peaks: p1 and p2) and VGlut 1 curves confirms a significant difference of the peak locations (**Fig. 3g**). Compared to the synaptic vesicle signal visualized by VGlut 1, which on average is homogeneously distributed throughout the presynaptic space, the peak α-tubulin (α-Tub) signal is located close to the outer synapse membrane. At least two peaks can be discriminated for F-actin: in close proximity to the outer and the inner presynaptic membrane. Part of the contribution to the second peak comes from the membrane-proximal part of the postsynaptic cell. Interestingly, βII-spectrin (βII-Spec) follows a similar distribution as F-actin and MyoVa is distributed similarly to VGlut 1.

Next, we investigated if protein distributions differed between line profiles drawn through AZ-containing calyx parts versus AZ-free areas, each reflecting an areas of specialized function (see **Supplementary Fig. 7**). Indeed, we encountered more VGlut 1 expression within a distance of up to 500 nm from the AZs (**Fig. 3h**), consistent with the distribution of SVs at sites of exocytosis known from electron microscopy^42^. Interestingly, less polymerized actin was found in close proximity to membranes containing AZs as compared to AZ-free membrane stretches (**Fig. 3i**). Strikingly, similar to VGlut 1, MyoVa was enriched at AZ-proximal regions, consistent with its role in SV trafficking (**Fig. 3j**). Exemplary plots of two individual AZs (**Fig. 3k, l**) demonstrate that different AZs can have differing fingerprints and further confirm a correlation between VGlut 1 and MyoVa.

Finally, to gain an overview of the global protein interrelationship, we conducted a large-scale colocalization analysis by determining Pearson’s r coefficients for all possible protein pairs at AZ-proximal versus AZ-free calyx patches (**Fig. 3m**). While synaptophysin 1 and VGlut 1 or different tubulin variants show expected colocalization with each other, irrespective of whether they are proximal to the AZ or not, other proteins differed significantly in their colocalization coefficient depending on their relation to the AZ. In agreement with the previous notion that less polymerized actin was found in AZ-proximity, F-actin showed significantly higher colocalization with WGA in AZ-free calyx regions. These results are consistent with previous work, suggesting that actin dynamics is linked to synaptic release^22,31,32,43^, yet for the first time directly visualizing this arrangement. Also the correlation between MyoVa and VGlut 1 already observed in the line profile analysis (**Fig. 3f**) can be confirmed by high colocalization coefficients, but is not significantly dependent on the proximity to AZs.

Based on our analyses of the overall (**Fig. 3f, g, Supplementary Fig. 8a-d**) and the AZ-dependent distribution (**Fig. 3h-m, Supplementary Fig. 8e-o**) of multiple targets at the MNTB synapse, we propose a model (**Fig. 4**) wherein F-actin and βII-spectrin are predominantly found at the calyceal borders with less F-actin at SV release sites. Tubulin density in AZ-containing calyx sections is higher in the center of the calyx lumen. SVs along with MyoVa are distributed throughout the calyx but show both increased levels in AZ proximity, suggesting that MyoVa does not only play a role in the initial transfer of SVs from microtubules to F-actin, but also in the later steps of the synaptic vesicle cycle, possibly including clustering and exocytosis. Although this model may appear familiar, it is the first of its kind that is derived from a set of 14 proteins identified at tens of nanometer resolution in the same physical sample, rather than reflecting a population average obtained from literally hundreds of different samples.

**Figure 4.**
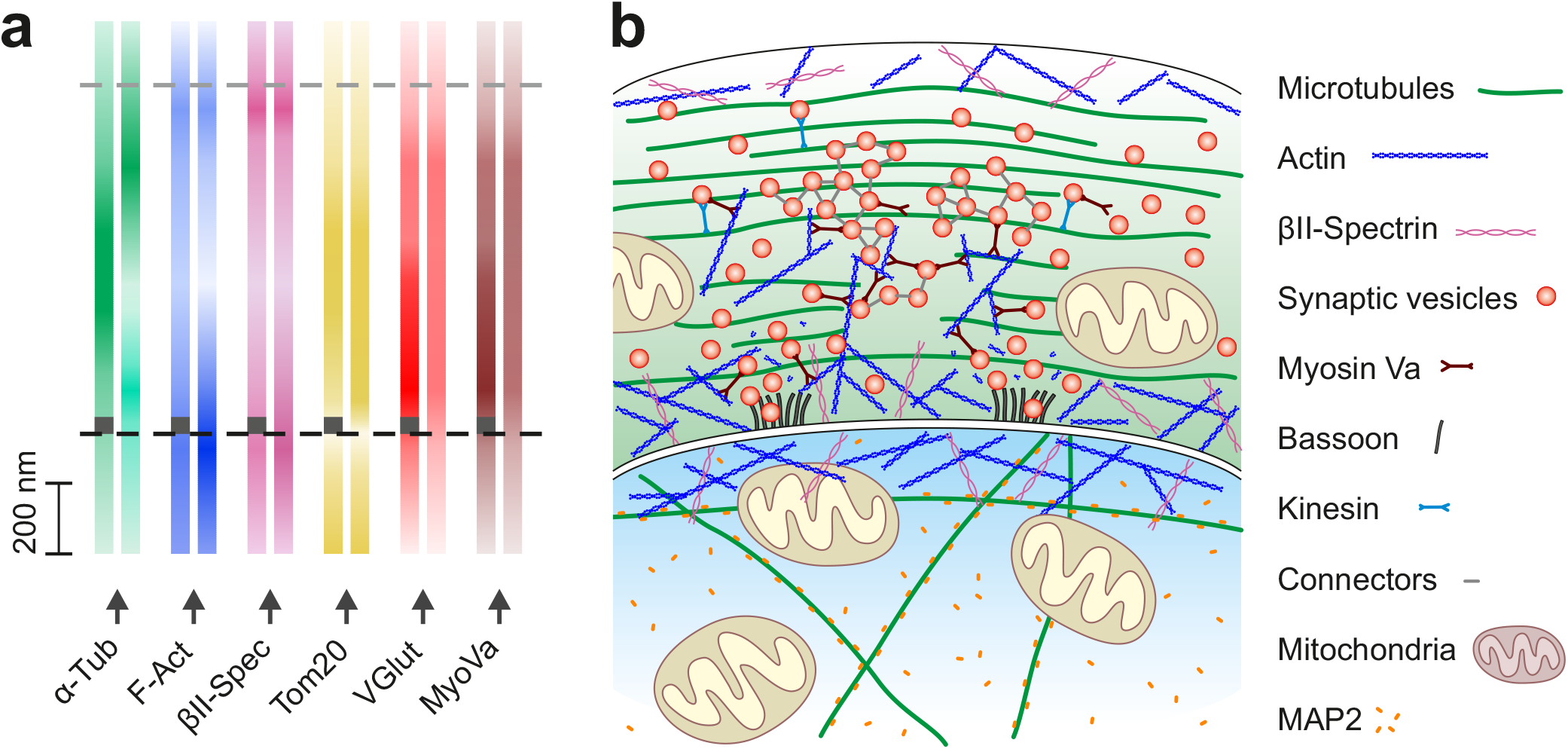
Model of protein distribution in the calyx of Held synapse. (**a**) Distribution of proteins examined in Figure 3 and Figure S8 indicated by color gradient. Distribution at active zone-positive regions are indicated by dark gray rectangles at the bottom of color gradient bars. (**b**) Model of synaptic architecture at the calyx of Held. Predominantly peripheral distribution of F-actin paralleled by βII-spectrin distribution. Note the decreased presence of polymerized actin in AZ regions. Microtubules are distributed throughout the presynapse with AZ-containing regions more densely populated by tubulins. Synaptic vesicles (SVs) cover up the whole presynaptic space. At the presynaptic membrane, SVs are more abundant in AZ-proximal regions. The distribution of myosin Va (MyoVa) is highly correlated with the vesicular VGlut 1 signal underpinning its role in SV trafficking: MyoVa is known to mediate the vesicular transfer from microtubules along which they are transported by kinesin motors to actin filaments. Interestingly, together with actin, MyoVa might be involved in SV clustering, similarly to connectors (structures of unknown molecular identity interconnecting SVs). In addition to the membrane-binding WGA lectin, the anatomy of the MNTB synapse was traced using bassoon as the AZ marker. While MAP2 is a protein that is specifically expressed at the postsynapse, the mitochondrial marker Tom20 is present in pre- and postsynaptic compartments, also confirming the geometry of this giant synapse.

## DISCUSSION

The maS^3^TORM strategy presented here successfully combines the advantages of three powerful approaches: super-resolution microscopy, multiplexing by optimized re-staining protocols, and automation. Careful consideration of label properties and thereof arising order of label application and removal techniques allowed us to optimize the number of targets detected in individual samples. We were able to automatically conduct high-content multiplexed SMLM imaging experiments in cells and neuronal tissue. Using our approach, we gained new insights on the multi-protein arrangement within single presynaptic terminals.

In contrast to spectral multiplexing, the maS^3^TORM approach avoids chromatic aberration by repeated usage of two fluorophores and spectral demixing. At the same time, we could drastically reduce the number of samples, which would be necessary to study protein interrelationships using the limited channel capacity of spectral multiplexing, and thus acquisition time. For example, being able to image only two targets at once, a researcher who wants to study 16 proteins, would have to conduct at least 120 single experiments without having the possibility to relate them to marker proteins such as WGA and bassoon required to clarify synaptic geometry. Furthermore, to conduct a comparable experiment and process the emerging data, researchers would need substantially longer working hours. By contrast, our approach minimizes the need of manual sample handling and ensures standardized conditions by automated liquid exchange. In addition, the looping functionality and modularity of the Experiment Editor enables an effective realization of any number of iterations. The feasible number of staining rounds is restricted mainly by the availability of labels with high specificity that allow efficient removal, and in case of labeling by primary and secondary antibodies by the limited number of antibody host species. Thus, broad availability of fluorophores directly conjugated to primary antibodies or other labeling agents will not only reduce the overall label size and thereby increase the spatial resolution, but also greatly improve multiplexing capacity. DNA-PAINT is another strategy that can easily be implemented in our automated microscope and allows for a high degree of multiplexing^7^. To further improve our computational efficiency, fiducial-based registration could be implemented into the post-processing software.

Deriving molecular organizational nano-architecture of subcellular compartments requires determining the position of as many proteins as possible within one and the same sample because some architectural features will only become evident when looking at more than two to three proteins and because averaging of pair-wise localization images across multiple samples may obstruct patterns that lack evident symmetries. An important prerequisite to achieve single probe multiplexing rests in the precision at which serial staining rounds can be superimposed, so that all proteins determined are placed in the same reference space. We demonstrate that this is possible with a precision of approximately 10-25 nm (**Fig. 3d**, see **Supplementary Note 2** for a detailed discussion), a number that could be further improved by using spatially invariant fiducial markers. Thus, maS^3^TORM offers a new avenue to systematically investigate complex nano-architectural principles of protein arrangement in single samples.

Finally, beyond demonstrating the functionality of maS^3^TORM, we also were able to gain new insights into the nano-architecture of a glutamatergic synapse. To achieve this, we established a tissue preparation method novel for super-resolution imaging: 400 nm thick tissue sections that are large enough to contain complete structures of interest while providing direct access of antibodies to the cytosolic compartment without the use of detergents. This approach ensures excellent structural preservation of the tissue and provides ideal conditions for super-resolution imaging. In such thin tissue sections, we observed more polymerized actin at AZ-free membranes as compared to AZ-proximal regions. This is in line with functional studies showing that actin dynamics regulates neuronal activity^22,30–32^. βII-spectrin shows a spatial distribution similar to actin at the calyx of Held and is decreased at presynaptic specializations, suggesting that actin and βII-spectrin may cooperate in synapse-modelling functions. Furthermore, we found that myosin Va distribution is correlated with VGlut 1 throughout the presynapse. In addition to its role in transferring SVs from microtubules to actin filaments^44^, this may imply additional functions of MyoVa: (1) interconnecting of SVs and/or (2) SV tethering to the membrane. Such connections could be realized via MyoVa-actin interactions^32^. The electron microscopic study by Cole et al. (2016)^45^ indicates that in addition to short SV connectors and tethers, longer cluster filaments are found either extending from the AZ or throughout SV clusters.

In conclusion, The maS^3^TORM approach proved to be an effective and universal method for studying highly complex biomolecular systems. With further development of labels, it will be possible to monitor more targets with even higher precision. Our data illustrates that elaborate computational methods employing artificial intelligence will be needed to infer organizational principles of protein nano-architectures from the distribution patterns of tens of proteins within an individual biological sample.

## METHODS

### maS^3^TORM setup

The custom-built 3D *d*STORM setup (**Supplementary Fig. 1a**, parts referenced here as #x and summarized in **Supplementary Table 4**) is equipped with 405 nm (#1; Cube, Coherent), 488 nm (#2; Obis, Coherent), 561 nm (#3; Obis, Coherent), and 661 nm (#4; Cube, Coherent) lasers. For excitation, lasers were focused onto the back focal plane of a 100× total internal reflection fluorescence objective with a numerical aperture of 1.49 (#9; Olympus). Emission light is collected by a multi-bandpass filter (#8; ZT405/488/561/647rpc, Chroma) and passes a cylindrical lens (#10; Thorlabs) that is used for astigmatism-based 3D imaging. Dual-color dSTORM imaging was performed by illumination with the 661 nm laser. Light emitted from the sample was filtered through the 700/75 ET bandpass filter, split into two beams by a custom-built dichroic filter (#13; 690 nm, Chroma), and focused onto the EMCCD camera (#15; iXon Ultra 897, Andor) with each channel occupying one half of the camera chip. The z-focus was stabilized by a feedback loop. The beam of an infrared diode (#16; 785 nm, Thorlabs) is reflected at the edge of the cover glass surface and the sample medium and projected onto a quadrant photodiode (#18; Laser Components). Depending on the position at which the beam illuminates the quadrant diode, a respective signal is sent to a motorized objective positioner (Physik Instrumente) to move the objective along the z-axis, thereby correcting for changes of the focal plane. To move the sample laterally, we used a custom-built motorized xy-stage (SmarAct). To enable an automated on-microscope staining and elution procedure, we complemented the microscope setup by the pipetting robot PAL3 RTC (Axel Semrau). Our robot is equipped with three vial trays and two syringes (100 μl and 1000 μl) and allows pre-mixing of solutions and their application onto the sample as well as their precise and reliable removal from the glass bottom dish.

For fully automated microscopy and re-staining procedure, both microscope hardware elements as well as the pipetting robot were integrated into one custom-written control software, called Experiment Editor that can be conveniently addressed by the user via the corresponding graphical user interface (GUI). To this end, all microscope components were implemented into the device manager of the open-source software μManager (Open Imaging)^35^. To conveniently use functionality of the hardware, initially, on top of μManager, the custom-written Microscope Control software was developed. The pipetting robot, in turn, can be controlled by the commercially available Chronos software (Axel Semrau). While the Microscope Control software can be directly addressed by the Experiment Editor, the pipetting robot is controlled and synchronized indirectly by employing a custom-written plugin that enabled Chronos to obtain tasks generated by the Experiment Editor in an exchange folder.

### Cell preparation

U2OS human bone osteosarcoma epithelial cells constitutively expressing Nup133-Ypet (Ypet is a YFP variant detectable by GFP nanobodies^46^) were cultured in DMEM/F-12 (Gibco) supplemented with 10% FCS (Gibco) and 2% GlutaMAX (Gibco) at 37°C in 5% CO_2_. Cells were transferred to glass bottom dishes (P35G-0-14-C, MatTek) pretreated with 50 μg/ml fibronectin (Sigma) and 1:500 Tetraspeck fluorescent beads (T7279, Invitrogen). After 24 h cells were washed twice with PSB and chemically fixed with 4% paraformaldehyde/4% sucrose dissolved in PBS for 15 min. After three washing steps with PBS, cells were permeabilized using 0.1% Triton in PBS for 10 min at room temperature (RT).

### Tissue preparation

All experiments on animals were conducted in compliance with the German animal welfare guidelines according to protocol G-75/15, and approved by the Regierungspraesidium Karlsruhe. Young (12 days old) Sprague-Dawley rats (Charles River) were anesthetized and transcardially perfused using ~20 ml PBS followed by ~30 ml 4% paraformaldehyde/PBS. Brains were removed and post-fixed in 4% paraformaldehyde/PBS for 24 h at 4°C. The brain stem was cut into 200 μm thick slices using a vibratome (SLICER HR2, Sigmann-Elektronik). Using a scalpel, the medial nucleus of the trapezoid body (MNTB) was cut out, resulting in a ~(200 μm)^3^ tissue block that was infiltrated in 2.1 M sucrose in 0.1 M cacodylic acid buffer (pH 7.4) for 1 h at RT. The tissue was placed on a specimen holder, plunge-frozen in liquid nitrogen, and further sliced into 400 nm sections using a cryo-ultramicrotome (Ultracut S with cryo-chamber EM FCS, Leica). Thin sections were picked up using a 2.3 M sucrose (in cacodylic acid buffer) droplet in a metal loop and transferred to glass bottom dishes that were previously incubated with 1:500 Tetraspeck beads. Prior to staining, tissue was thawed for 10 min at RT and sucrose was removed by three washing steps with PBS for 15 min. This method of tissue preparation ensures direct access of antibodies to epitopes within the cell without the use of detergents.

### Multiplex procedure

#### Experiment Editor

All steps described below were carried out using the custom-written Experiment Editor software (**Supplementary Fig. 1b**). Its graphical user interface (GUI) allows convenient and coordinated operation of microscope components and the pipetting robot (**Supplementary Fig. 2**). Microscope control was realized via custom-written Microscope Control software, which, in turn, addresses μManager, an open source software, where all microscope hardware components were integrated. The pipetting robot was operated using a handshake procedure communicating via a specific exchange folder. A plugin that was custom-written for the commercial Chronos software constantly checks the content of an exchange folder for a path to a Chronos-tailored task list generated by Experiment Editor. The task list consists of a set of commands that is loaded into Chronos and executed by the robot. After the tasks are successfully completed, a text file with specific key words is saved to the exchange folder by the Chronos plugin. This file, in turn, instructs the Experiment Editor to continue with the execution of the next workflow module.

All staining, bleaching and elution procedures were carried out by the pipetting robot at room temperature (21 °C).

#### Staining procedure

Cells and cryo-sections were both blocked in 5% fetal calf serum (FCS) for 10 min. For immunohistochemistry, antibodies were applied in 0.5% FCS for 1 h. After each application of antibodies or other labels, samples were washed three times with PBS for a total of 15 min. Alexa Fluor 647 conjugated nanobodies against GFP and CHC17 against clathrin heavy chain 1 were applied for 1 h in 0.5% FCS. Alexa Fluor 647 conjugated phalloidin and CF680 conjugated wheat germ agglutinin (WGA) were applied in PBS for 20 min. JF_646_-Hoechst was directly added to the imaging buffer. For the precise Re-STORM workflow used for U2OS cells (**Fig. 2a**) and for MNTB slices (**Fig. 3c**), see **Supplementary Tables 1** and **2**, respectively. All antibodies and other staining reagents used for cells and neuronal tissue are summarized in **Supplementary Table 3**.

#### SMLM image acquisition

For all imaging rounds, β-mercaptoethylamine (MEA) buffer at pH 8 containing 100 mM MEA and 15 mM KOH in PBS was freshly prepared by the pipetting robot. To prevent dilution of MEA buffer by residual liquid inside the glass bottom dish, before imaging, the sample was once washed with MEA buffer. Conventional wide field images were acquired using 661 nm laser at low power (~1 W/cm^2^) with an exposure time of 200 ms. For *d*STORM images, photoswitching of fluorophores was initiated by 10 s exposure to 661 nm laser at high power (~2 kW/cm^2^) followed by the acquisition of 20,000 frames at an exposure time of 30 ms. PAINT imaging with JF_646_-Hoechst was performed at an exposure time of 50 ms. All images were recorded in astigmatism-based 3D mode and automatically, via a Python script, redirected to and processed by rapidSTORM.

#### Fiducial imaging

For precise registration of images from distinct staining rounds, we imaged fluorescent bead fiducials (excitable at 560 nm, 660 nm, and other wavelength; Tetraspeck beads T7279, Invitrogen). To ensure maximal immobilization, we seeded the fiducials onto glass bottom dishes underneath the cells or tissue. As STORM images are performed at very high 661 nm laser intensity, bead fluorophores with an excitation maximum of 660 nm are bleached after approximately two STORM and two bleaching rounds. To preserve fiducials for more than ten imaging and bleaching rounds, we used weak excitation by the 561 nm laser. Recording through a filter (700/75 ET Bandpass, AHF: F47-700) optimized for the 661 nm laser allowed us to minimize chromatic aberration. Regions of interest (ROIs) for SMLM imaging were chosen such that they contained at least five fiducials to allow precise registration.

#### Auto-refocusing system

Automated focus recovery was implemented based on Tetraspeck fiducials. A reference region with at least five fluorescent beads in focus (without cells or tissue) was selected and its coordinates were transferred to the automated refocusing module. Subsequently, the major multiplex workflow was launched. In our design applied for cells and nervous tissue, initially the bead-reference region is approached, images are taken, automatically processed by rapidSTORM and the mean z-values for the beads are extracted. After that, the Auto Focus module moves the objective through six focus positions (customized step size); for each position the mean z-value is determined. Finally, the z-position with a z-value closest to the reference z-value is selected. Next, STORM images of the preselected ROIs (five in our experiment design) are acquired and automatically preprocessed by rapidSTORM.

#### Chemical elution

In cells and neuronal tissue, elution of antibodies was realized using 0.1% SDS at pH 13 for 15 min (modified from Collman et al. (2015)^37^. To ensure the desired concentration, the sample was once washed with elution buffer before it was applied for a second time for incubation. After elution, the sample was washed with PBS six times in total.

#### Photo-bleaching

Bleaching was performed in PBS using the 661 nm laser at 100% power (~2 kW/cm^2^) and 405 nm laser at 50% power (0.7 kW/cm) for 4 min ^10^.

### Series of steps in an automated imaging session

Each experiment consists of a series of steps illustrated in **Figure 1c**. After preparing and fixing the sample (1), the experimental design is programmed using the custom-written Experiment Editor (2), the sample is stained for the first time (staining round 1) either manually or semi-automatically using the pipetting robot (3), and vials with solutions for labeling, washing and elution are prepared. Subsequently, regions of interest (ROIs) are selected for acquisition (4). To prevent focus loss during a long-term experiment, an automated refocussing device maintains the focal plane based on a preselected region containing fluorescent bead fiducials (5). Then *d*STORM 3D images of two channels are acquired and spectrally demixed (6). The new round starts with the acquisition of bead fiducials for subsequent registration of images from different staining rounds (7). To remove labeling from the imaged staining round, depending on the type of labels used, chemical elution (8) and/or photobleaching (9) is performed. Thereafter, the sample is re-stained using a new set of labels. (10) Refocusing (11) precedes dSTORM 3D image acquisition. Finally, the procedure can be repeated as many times as desired by resuming with step 7. After a completed multiplex session, data sets were individually evaluated regarding labeling and image quality, and high-quality SMLM images were qualified for further analysis.

### Imaging experiments on U2OS cells

U2OS cells constitutively expressing the nuclear pore component Nup133 tagged with Ypet were successively stained (**Fig. 2a, Supplementary Table 1**) with Alexa Fluor 647 conjugated nanobodies against GFP that bind Ypet (SR1/Ch1), phalloidin binding actin (SR2/Ch1), WGA-CF680 binding sialic acid and N-acetylglucosaminyl sugar residues at the plasma membrane, at the Golgi trans face and in the center of nuclear pores (SR2/Ch2), Alexa Fluor 647 conjugated clathrin antibodies (SR3/Ch1), anti-tubulin (SR4/Ch1), anti-paxillin (SR4/Ch2), anti-vimentin (SR5/Ch1), cis-Golgi recognizing anti-GM130 (SR5/Ch2), anti-fibrillarin (SR6/Ch2), mitochondria-specific anti-Tom20 (SR7/Ch1), anti-lamin A/C (SR8/Ch1), ER recognizing anti-PDI (SR8/Ch2) antibodies, as well as early endosome anti-Rab5 (SR9/Ch2) and anti-EEA1 (SR10/Ch1) antibodies, and a Hoechst-JF646 conjugate that reversibly binds DNA (SR11/Ch1). In addition to this representative experiment, another three experiments were carried out with similar designs and outcomes.

### Imaging experiments at the calyx of Held giant synapse

Tissue was labeled with antibodies against bassoon (SR1/Ch1) and homer 1/2/3 (SR1/Ch2), phalloidin (SR2/Ch1) and WGA-CF680 (SR2/Ch2), fluorophore-labeled CHC17 primary antibody (SR3/Ch1), and antibodies against MAP2 (SR4/Ch1) VGlut 1 SR4/Ch2), GM130 (SR5/Ch2), Rab3a (SR6/Ch1), Tom20 (SR6/Ch2), α-tubulin (SR7/Ch2), γ-actin (SR8/Ch1), synaptophsin 1 (SR8/Ch2), α/βsynuclein (SR9/Ch1), PDI (SR9/Ch2), and Hoechst-JF646 conjugate that reversibly binds DNA (SR10/Ch1) (**Fig. 3c, Supplementary Table 2**). In addition to this representative experiment, another six experiments were carried out with similar designs and outcomes.

### SMLM image reconstruction and registration

Directly after SMLM acquisition, employing a Python script, image frames were automatically redirected to rapidSTORM software for single-molecule localization and image reconstruction. For subsequent SMLM processing steps, including spectral demixing, drift correction, and 2D as well as 3D rendering, the Post-Processing software was developed. To allow custom data processing, we complemented the software by a GUI with modular design. Fast and standardized image analysis was possible using the batch processing function. SMLM images from consecutive staining rounds were registered based on fluorescent bead fiducials (at least five). SMLM images were linearly transformed based on mean distances along x- and y-axes between corresponding fiducials from different imaging rounds. For the registration error, the average difference between the distance of single beads and the mean distance of all beads was calculated.

### Analysis of protein distribution at the calyx synapse

The protein distribution within the calyx of Held was analyzed using discrete line profiles manually drawn in ImageJ (**Supplementary Fig. 7**). Initially, the tightly apposed pre- and postsynaptic membranes labeled by WGA were traced by the “segmented line selection tool”. Using a custom-written ImageJ macro, the line was automatically interpolated, split into 1 μm segments, and rotated by a 90° angle. This ensured an orientation of line profiles perpendicular to the synaptic membrane. The line selections were elongated and the line thickness was set to 700 nm. To ensure reliable analysis, we considered contiguous membranes, which we defined as a clearly discernible WGA staining at both membranes (outer and inner) of the synapse that is traceable in at least 70% of the analyzed selection. Correct identification of synaptic membranes was additionally confirmed by AZ and SV markers. For all calyx segments that passed our criteria, multi-line plots were automatically generated and saved as tables. Data tables from 5 experiments (with 3 calyces per experiment and 4-16 line profiles per calyx) were collected and imported to Matlab for further analysis. We define the outer and inner (synaptic) borders of the calyx as presynaptic membrane not in touch with the postsynaptic cell and presynaptic membrane touching the postsynaptic cell, respectively. The pre- and postsynaptic membranes constituting the inner border cannot be discriminated because they are separated by less than 20 nm in aldehyde-fixed tissue (unpublished electron microscopic measurements). The calyx borders were approximated by fitting the line profiles with the Gaussian function. The mean distance between the outer and the inner calyx border was calculated from the whole WGA data set. All line profiles of the WGA signal and of all other proteins analyzed were scaled and aligned according to the borders of an average calyx. The difference between the calyceal distribution of α-tubulin, F-actin and VGlut 1 was quantified by localizing their intensity peaks using the Gaussian function. Statistical significance was evaluated by one-way ANOVA followed by Bonferroni post-hoc test.

For the analysis of AZ-positive versus AZ-negative calyx segments, 300 nm thick line profiles were laid across two AZ-containing calyx regions (characterized by dense Bassoon signal) and two regions free of AZs per calyx (**Supplementary Fig. 7b, c**). In total, 13 calyces from 5 independent experiments were used for this analysis. The same dataset was used for the colocalization analysis. In this case, line profiles were manually restricted to the calyx and transformed to area selections. Correlation between all possible combinations of proteins within these selections was automatically (ImageJ macro) determined using the Pearson’s coefficient. Data was exported to Matlab and extracted to colocalization matrices, which in turn have been further processed, averaged and compiled as a heatmap in Excel. Colocalization data of selected pairs of proteins in AZ-positive versus AZ-negative regions were statistically analyzed by unpaired two-tailed Student’s t-test

## Supporting information

Supplementary Information index

## ACKNOWLEDGMENTS

The purchase of the PAL3 pipetting robot was supported by the CellNetworks Excellence Cluster (EXC 81). We thank Heinz Horstmann for training M.K. in cryo-sectioning of MNTB tissue, Claudia Kocksch for excellent technical assistance. We also thank M. Simonetti (R. Kuner laboratory) for the Vimentin antibody, C. Spahn for the CHC17 antibody, and Luke Lavis for JF_646_-Hoechst.

## AUTHOR CONTRIBUTIONS

M.K. designed and performed all experiments, developed computational image analysis tools, analyzed all data and generated all images. F.H. with assistance of M.K. built the maS^3^TORM setup and developed the acquisition interface “Experiment Editor”. F.H. designed and developed the Post-Processing software for STORM images. V.V. devised the concept of automating the STORM setup, conducted preliminary tests with a demo version of the robot and pretested elution and bleaching protocols. S.S. performed control experiments and produced Supplementary Video 1. M.H. and T.K. contributed to experimental design, data analysis, and supervised this work. M.K., M.H., and T.K. wrote the manuscript. All authors edited and approved the manuscript.

## Data availability

All raw and processed data will be made available upon request.

## Code availability

Microscope Control software with integrated Experiment Editor software (custom code: https://github.com/fherrmannsdoerfer/MicroscopyControl2), Chronos 4.8.0 (commercial). Plugin for Chronos (custom code: https://github.com/fherrmannsdoerfer/Chronos_Plugin).

