## Supplementary Information index for "Automated highly multiplexed super-resolution imaging of protein nano-architecture in cells and tissues"

**Supplementary information guide**

| **Supplementary Element** | **Title** | **Location** |
| --- | --- | --- |
| Supplementary Figure 1 with legend | Microscope components and software used to control the maS^3^TORM setup. | SI file, pp. 1-2 |
| Supplementary Figure 2 with legend | Graphical user interface (GUI) of the Experiment Editor software. | SI file, pp. 3-4 |
| Supplementary Figure 3 with legend | Control experiments for bleaching- and elution-mediated signal removal. | SI file, pp. 5-6 |
| Supplementary Figure 4 with legend | Single case control experiments for estimation of cross-talk and labeling efficiency. | SI file, pp. 7-8 |
| Supplementary Figure 5 with legend | Localization precision analysis for images acquired throughout the multiplex experiment. | SI file, p. 9 |
| Supplementary Figure 6 with legend | Examples showing 3D visualization of selected STORM images. | SI file, p. 10 |
| Supplementary Figure 7 with legend | Workflow for analysis of presynaptic architecture of the calyx of Held. | SI file, p. 11-12 |
| Supplementary Figure 8 with legend | Averaged line profiles of global and active zone-specific protein distributions. | SI file, p. 13 |
| Supplementary Figure 9 with legend | Colocalization matrix. Pearson’s r values for the colocalization matrix shown in Figure 3m. | SI file, p. 14 |
| Supplementary Note 1 | Extended information for control experiments for signal removal (Supplementary Fig. 3a) | SI file, pp. 15-16 |
| Supplementary Note 2 | Supplementary discussion of registration precision between different staining rounds for cells or tissue | SI file, p. 16 |
| Supplementary Table 1 | Experimental workflow for multiplex experiment in U2OS cells shown in Figure 2a. | online |
| Supplementary Table 2 | Experimental workflow for the multiplex experiment in the medial nucleus of the trapezoid body shown in Figure 3c. | online |
| Supplementary Table 3 | All antibodies and other labels used in this work. | online |
| Supplementary Table 4 | maS^3^TORM components. | online |
| Supplementary Table 5 | Exact number of experiments, samples, and selections for all quantified analyses. | online |
| Supplementary Video 1 | Performance of the maS^3^TORM setup. | online |
| Supplementary Video 2 | Super-resolution images of three target proteins from different imaging rounds merged in 3D space. | online |

SI = Supplementary Information

**Supplementary Video 1 | Performance of the maS^3^TORM setup.** Video demonstrating buffer preparation, liquid exchange, approaching sample regions and image acquisition by maS^3^TORM.

**Supplementary Video 2 | Super-resolution images of three target proteins from different imaging rounds merged in 3D space.** STORM images of phalloidin (red), α-tubulin (green), and Tom20 (blue) rendered and merged in 3D.
