## Supplementary Information for "Automated highly multiplexed super-resolution imaging of protein nano-architecture in cells and tissues"

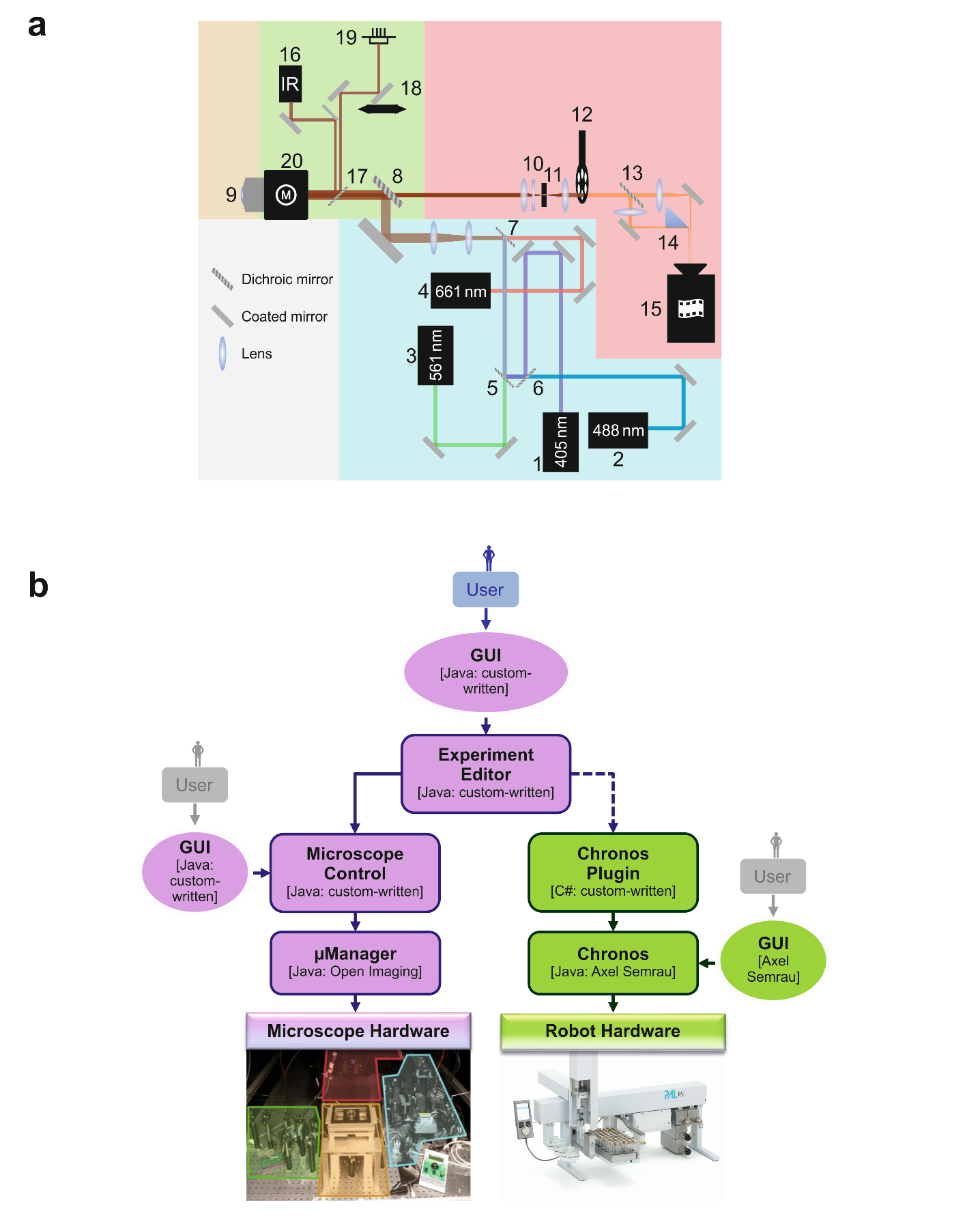

**Supplementary Figure 1 |** **Microscope components and software used to control the maS^3^TORM setup.** (**a**) Detailed schematic of the microscope components of the maS^3^TORM setup (see also **Supplementary Table 4**). For emission (blue background), the home-built *d*STORM system is equipped with four lasers: 405 nm (#1), 488 nm (#2), 561 nm (#3), and 661 nm (#4). The 405 nm, 488 nm, and the 561 nm laser beams are merged using two dichroic mirrors (#5 and #6). The 661 nm laser beam that is primarily used for *d*STORM measurements is conjoined with all the other laser beams by the dichroic mirror (#7). Subsequently, the excitation light is widened by two lenses forming a telescope and is reflected onto the sample using another dichroic mirror (#8). The emission beam (red background) is collected by the objective (#9; yellow background) and passes a tube lens followed by the cylindrical lens (#10) that can be used for 3D imaging. At the focal point of the tube lens a slit (#11) is placed, followed by another lens parallelizing the beam. Different emission filters to block excitation light are mounted in a filter wheel (#12). The dichroic filter (#13) splits the emission light into two beams with one beam passing and the other being reflected at the inner edge of a prism (#14) resulting in two channels that are focused onto the camera (#15). The microscope is further provided with an infrared beam-based active focus stabilization system (green background). The incoming beam of the infrared diode (#16) is coupled into the objective by the dichroic mirror (#17). The beam is reflected at the edge between the glass surface of the sample and the sample medium. Using a movable mirror (#18), the reflected beam is projected onto a quadrant diode (#19). To correct for focus drift, the quadrant diode is linked to a controller (not shown) that, in turn, initiates movement of a piezo-driven stage (#20) wherein the objective is mounted. (**b**) Software architecture designed for coordinated control of *d*STORM and robot hardware components. All microscope hardware components (photograph in the left lower part of the diagram) were implemented into the open-source software called µManager^1^. On top of µManager, the custom-written Microscope Control software and the corresponding graphical user interface (GUI) were developed to conveniently use all necessary functionality of the hardware. The setup was complemented by the commercial pipetting robot PAL3 by CTC Analytics AG/Axel Semrau (photograph in the right lower part of the diagram) and can be controlled by the commercial Chronos software (Axel Semrau). For a fully automated microscopy and re-staining procedure, both microscope hardware elements and the pipetting robot were integrated into one custom-written Experiment Editor software that can be conveniently addressed by the user via the corresponding GUI depicted at the top center of the diagram. While the Microscope Control software is directly addressed by the Experiment Editor software, the communication between Experiment Editor and the pipetting robot is realized indirectly (depicted by the dashed line) via an exchange folder that is constantly checked and modified by the custom-written plugin for Chronos as well as by the Experiment Editor.

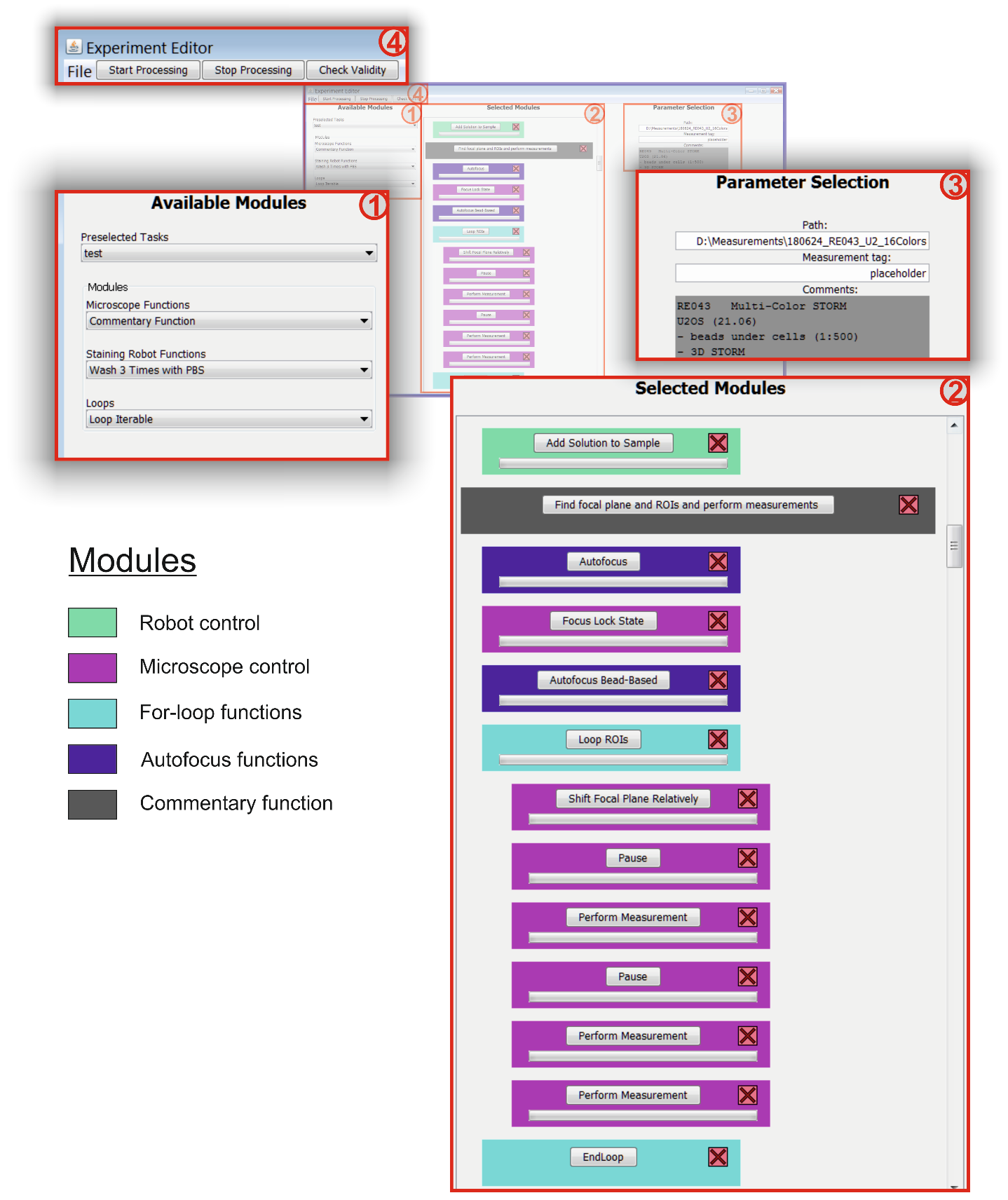

**Supplementary Figure 2 |** **Graphical user interface (GUI) of the Experiment Editor software.** Zoom-in into main Experiment Editor GUI parts. To create a customized experiment, the user can select modules from three main categories listed in the ‘Available Module’ section (1). The ‘Microscope Functions’ dropdown list contains all modules that directly control the hardware of the microscope such as lasers, stage, or camera. The ‘Staining Robot Functions’ list contains modules that trigger different actions of the robot. Most modules bundle different robot actions together for a convenient and time-saving usage. The ‘Loops’ dropdown category contains two loop modules: ‘Loop Iterable’ and ‘Loop ROIs’. ‘Loop Iterable’ is used to repeat modules of choice for a desired number of times while ‘Loop ROIs’ is designed to repeat microscopic actions for a list of chosen ROIs that are automatically approached by the microscope stage. Once a module is chosen from one of the three categories, it appears in the ‘Selected Modules’ column in the center of the GUI (2). An exemplary excerpt of a typical experiment is shown. It contains all types of modules: robot control-related (green), microscope control-related (magenta), and loop functions (light blue). In addition, autofocus functions (dark blue) and the commentary function (gray) can be selected from the ‘Microscope Control’ list. After being added to the ‘Selected Modules’ field, the order of all modules can be changed by the drag and drop principle. By clicking the cross on red background, modules can be deleted from the list. Gray push-buttons of individual modules can be clicked to display and modify module-specific parameters, which are shown in the right ‘Parameter Selection’ part of the GUI (3). After an experiment is designed, its consistency can be checked by the ‘Check Validity’ button on the upper left corner of the GUI (4). This function performs simple checks to ensure that, for example, each parameter field is filled and that loops have the right number of parameters corresponding to the number of repetitions. Subsequently, the workflow can be executed by clicking the ‘Start Processing’ and interrupted by the ‘Stop Processing’ button. The ‘File’ menu provides possibilities to save and load workflows.

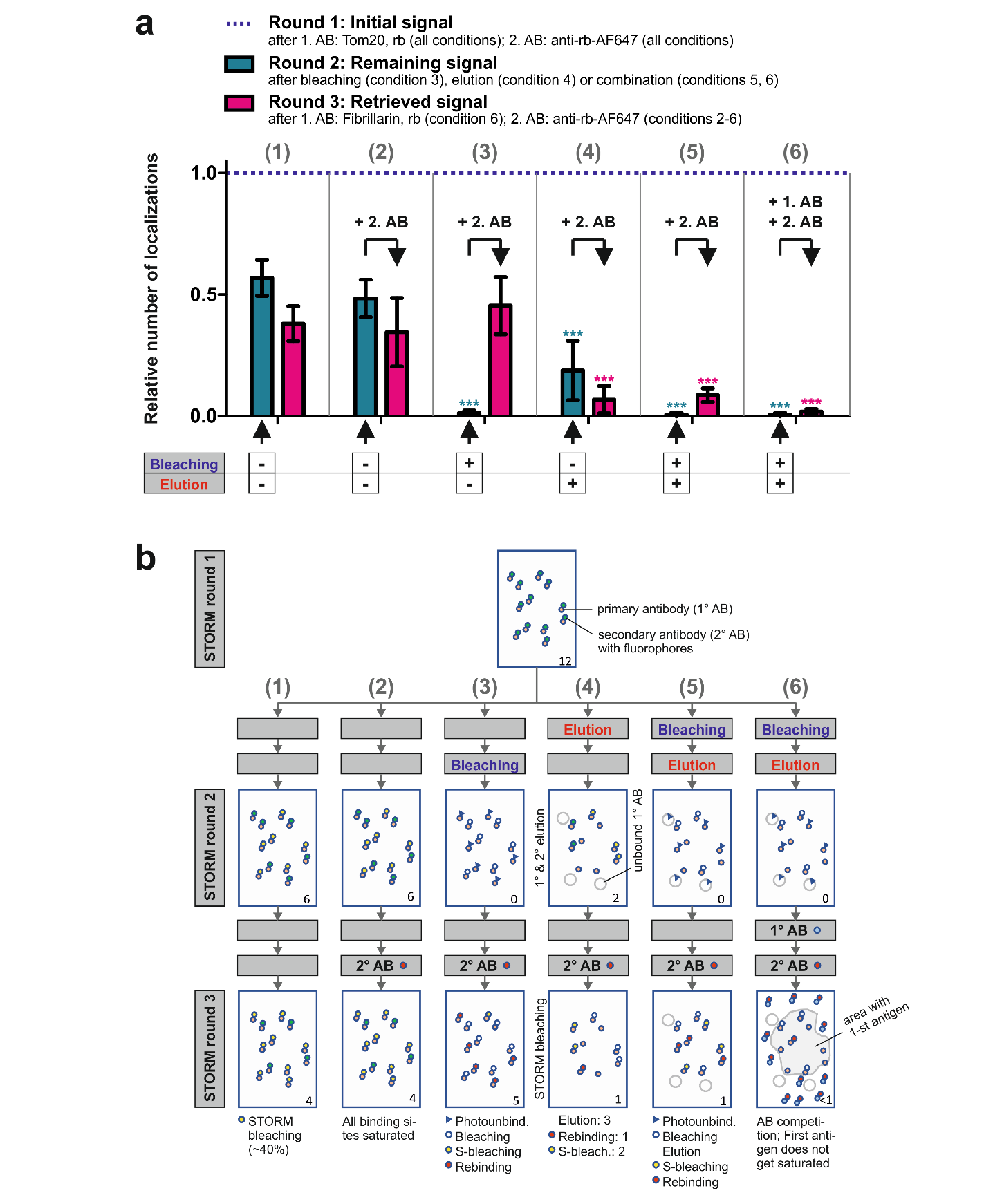

**Supplementary Figure 3 |** **Control experiments for bleaching- and elution-mediated signal removal.** (**a**) Six test experiments were carried out to evaluate the cross-talk from consecutive multiplexed STORM rounds. For all six test conditions, U2OS cells were stained with a primary antibody against Tom20 from rabbit (rb) and a secondary anti-rabbit antibody conjugated to the Alexa Fluor 647 (AF647) fluorophores. Three STORM imaging rounds were carried out for each condition. Signal intensity from the initial staining acquired in the first imaging round was evaluated by quantifying localizations within selections of Tom20-positive regions. Signal from the following two rounds was normalized to the first round. In the second imaging round, the influence of elution and/or bleaching was evaluated. Bleaching was either induced by the STORM acquisition itself or by the intended photobleaching procedure (conditions described in the Methods section). Prior to the third imaging round, for test conditions 2-6, the secondary antibody against rabbit was reapplied and the percentage of signal retrieval was evaluated. To measure the influence of a competing primary antibody, which is usually present during a typical multiplex experiment, in test condition 6, we also applied an antibody from rabbit against Fibrillarin as a test case. This condition resulted in a cross-talk of ~1.8%. For a detailed description of the six test conditions, see **Supplementary Note 1**. Each experiment was carried out for 3 independent sample dishes with 3 cells analyzed per sample dish. For Tom20 signal quantifications, 3 selections in mitochondria-positive area were analyzed per cell. Background signal was determined in cell area lacking mitochondria (3 selections/cell) and subtracted from Tom20 signal to quantify specific Tom20 signal. Bars represent mean ± SD. For statistical analysis, all bars corresponding to imaging round 2 were compared to round 2 of the first test condition. In analogy, all bars corresponding to imaging round 3 were compared to round 3 from the first test condition. Asterisks indicate statistical significance (*** corresponding to p < 0.001), assessed by one-way ANOVA followed by Bonferroni post-hoc test. (**b**) Illustration of effects measured in **Supplementary Figure 3a**. Exemplary numbers of antigens labeled by antibodies with intact fluorophores are shown in the right bottom corner of each panel illustrating an imaging round.

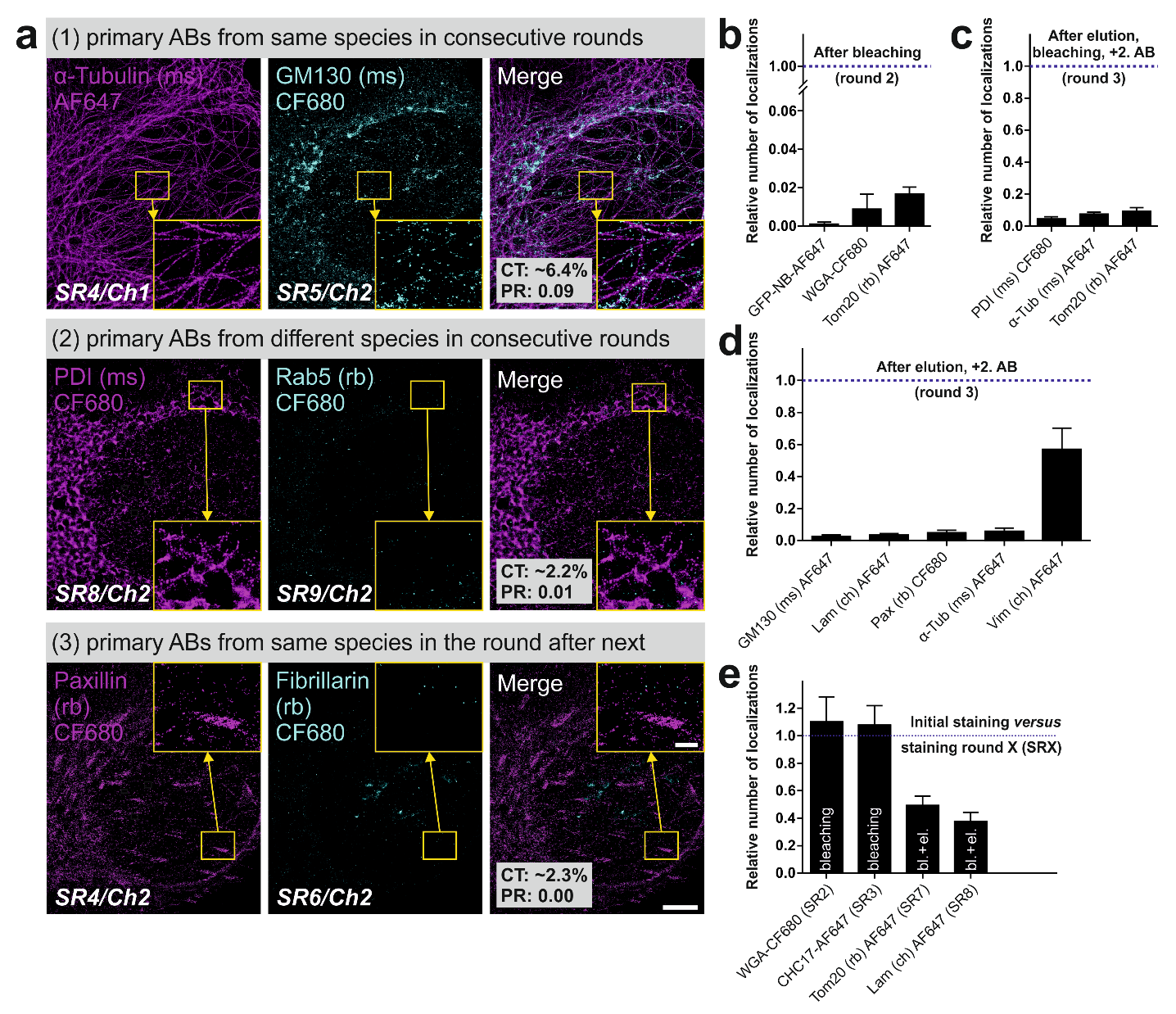

**Supplementary Figure 4 |** **Single case control experiments for estimation of cross-talk and labeling efficiency.** (**a**) Three examples from the multiplex experiment in **Figure 2a** were analyzed with respect to unwanted cross-talk from preceding staining rounds (SR). Here, after each SR elution and bleaching procedures were carried out. In the first test case (upper row of panels) images from consecutive rounds were analyzed where primary antibodies from the same species, in this case mouse (ms), were used. The initial α-tubulin staining resulted in a cross-talk (CT) of ~6.4% in the GM130 channel (Ch) of the consecutive imaging round. Colocalization analysis also resulted in a relatively low Pearson’s r (PR) coefficient of 0.09. Only ~2.2% CT and 0.01 PR were estimated for primary antibodies from different species used in consecutive rounds (central panel row). Whenever possible, primary antibodies from the same species were not used in the consecutive round but in the round after the next. Such a case exemplified by Paxillin staining in round 4 and Fibrillarin staining in round 6 resulted in CT ~ 2.3% and PR = 0 (lower panel row). Scale bars correspond to 5 µm in main panels and 1 µm in magnified boxed areas. (**b**) The effect of bleaching in consecutive rounds was assessed based on three examples yielding very low cross-talk values of ~0.1% to ~1.7%. (**c**) The combination of elution and bleaching with subsequent reapplication of the same secondary antibody (2˚ AB) in the next round resulted in cross-talk values ranging from 4.5% to 9.3%. (**d**) The effect of elution (without the additional bleaching step) was evaluated based on five examples resulting in values below 6% for most test cases. Unexpectedly, the vimentin antibody showed an extraordinarily high affinity and resulted in a cross-talk of ~56.9%. Therefore, in our experiments we did not use other antibodies from chicken in consecutive rounds. (**e**) To assess signal quality with increasing numbers of rounds after bleaching or bleaching and elution, we compared the labeling efficiency of four test antibodies from the multiplex experiment in **Figure 2a** to their performance in fresh untreated cells (round 1). After one or two bleaching rounds, labels (in this case WGA lectin and CHC17 antibody) were as efficient as on untreated cells. Despite numerous elution and bleaching rounds, Tom20 antibody application in SR7 still resulted in ~49.2% of the initial signal; and ~37.6% of initial performance could be reached by the lamin antibody in SR8. (**b**-**e**) For sample numbers, see **Supplementary Table 5**. Bars represent mean ± SD. Abbreviations: ms = mouse, rb = rabbit, ch = chicken, AF647 = Alexa Fluor 647, bl. = bleaching, el. = elution.

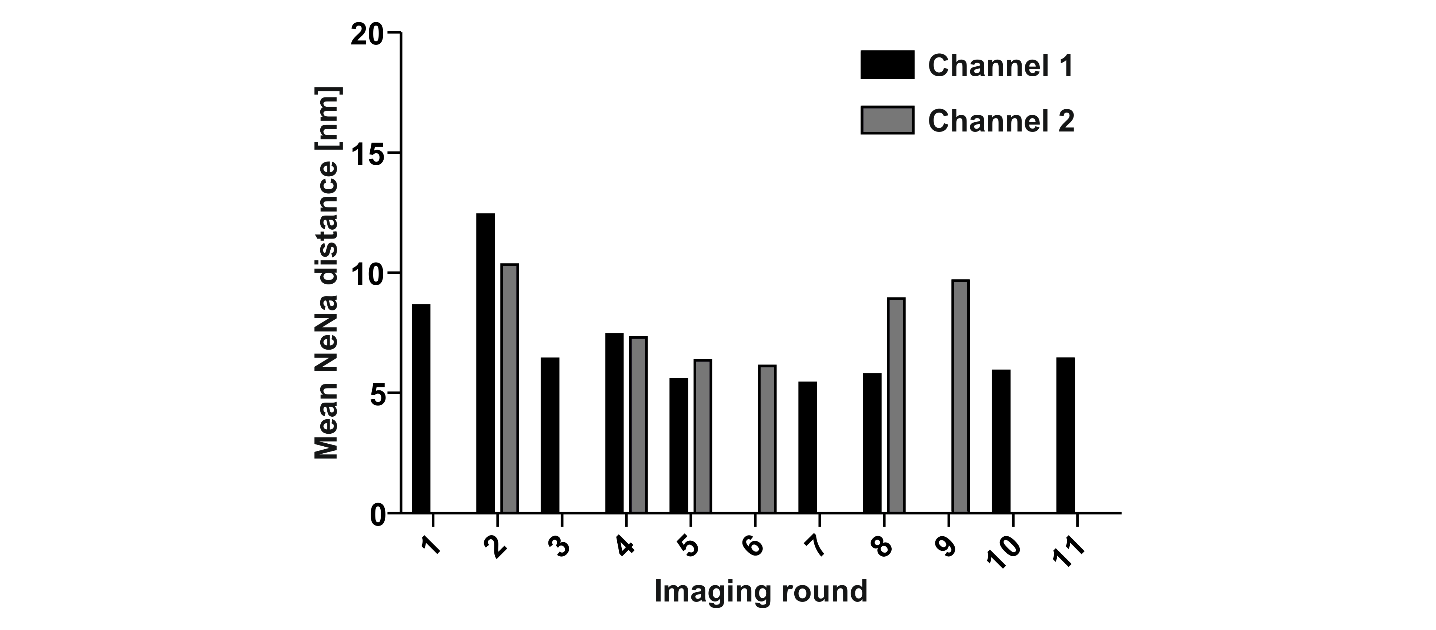

**Supplementary Figure 5 |** **Localization precision analysis for images acquired throughout the multiplex experiment.** Localization precision was estimated by quantifying the nearest neighbor (NeNa) distance between corresponding localizations from adjacent STORM frames^2^. NeNa analysis resulted in a mean NeNa distance of 7.7 ± 2.7 for all imaging rounds, indicating a very high localization precision. For this analysis we used the NeNa algorithm of LAMA software^3^.

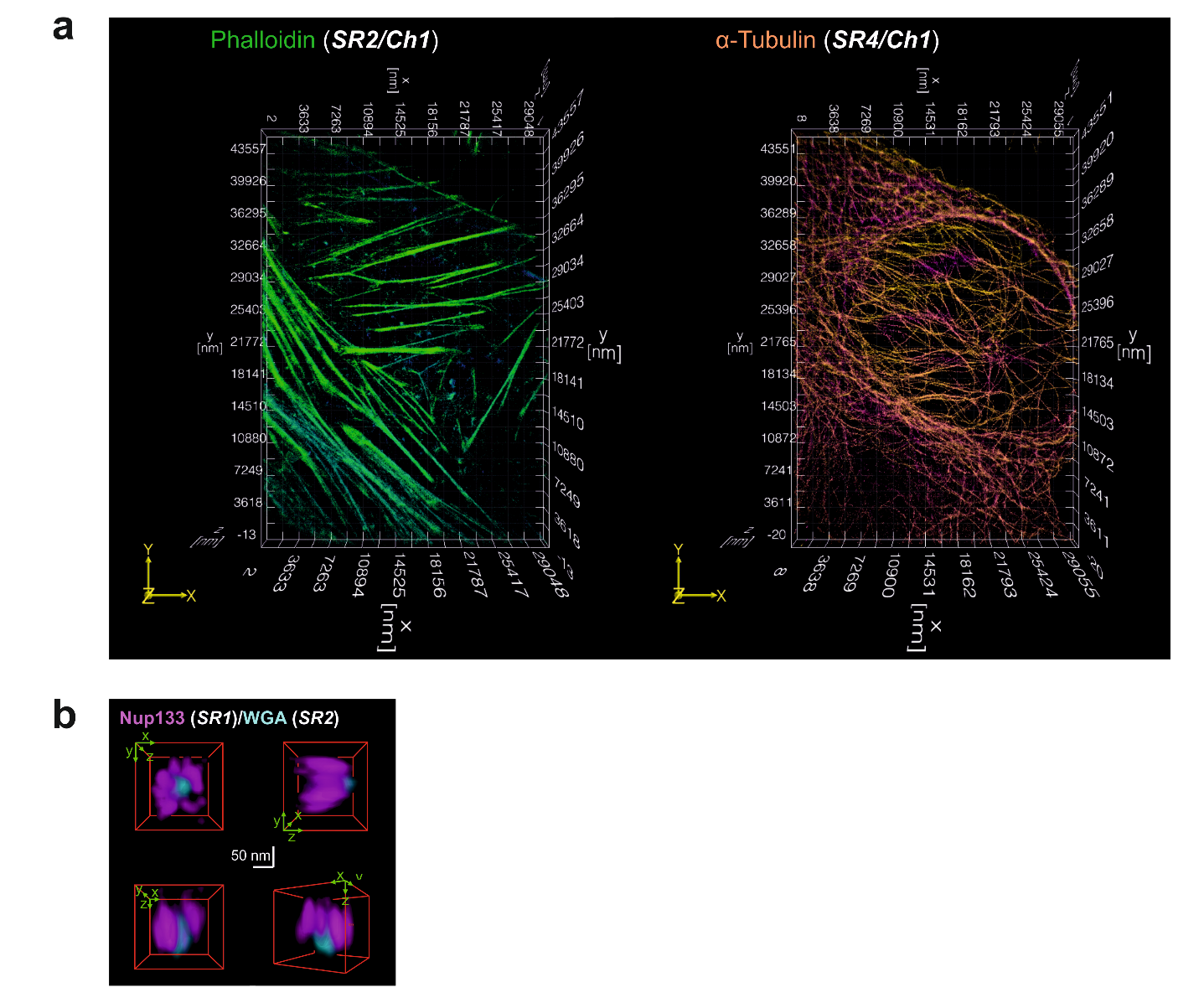

**Supplementary Figure 6 |** **Examples showing 3D visualization of selected STORM images.** (**a**) Three-dimensional representation of the phalloidin staining (left panel) and α-tubulin staining (right panel) from the multiplex experiment shown in **Figure 2a** using ViSP software^4^. (**b**) Second example (for first example, see **Figure 2d**) of a nuclear pore complex rendered in 3D, demonstrating high axial resolution and robust fiducial-based registration precision. ImageJ was used for 3D visualization in **b**.

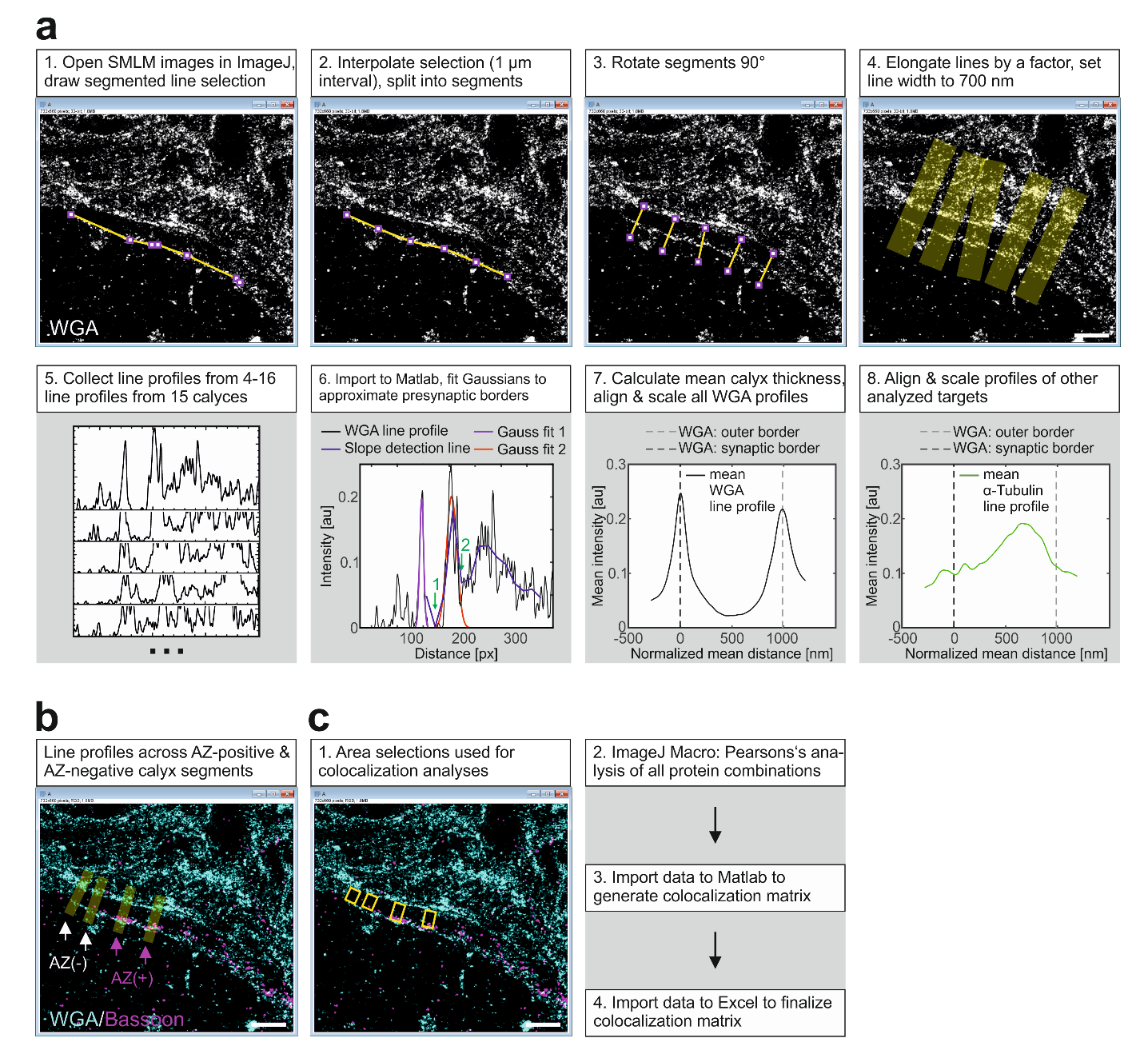

**Supplementary Figure 7 | Workflow for analysis of presynaptic architecture of the calyx of Held.** (**a**) The distribution of presynaptic proteins was analyzed using a custom-written plugin for ImageJ^5^. In the first step (1), the user has to roughly outline the inner border of the calyx by a segmented line selection. Outlining was performed based on images with WGA staining aided by information from stainings of active zone (AZ) marker Bassoon and synaptic vesicle markers (e.g. VGlut). Subsequently, the plugin can be launched and the line selection is automatically interpolated and split into segments (2). The segments are rotated 90° (3) to ensure that line profiles will be determined perpendicularly to the inner calyx membrane. In the next step (4), lines are elongated to cover the whole presynapse and their width is set to a desired value (for this analysis to 700 nm). The ‘multi plot’ command is executed to collect line profiles from all line profiles per calyx. Line profiles from all analyzed calyces are collected as text files (5) and imported to Matlab for further analysis. Peaks corresponding to calyceal borders are fitted by Gaussian functions (6). For this purpose, the inner calyx border is localized first within a defined window; this fit is very reliable as the distance between the line profile origin and the inner border is nearly constant. As different calyces and calyx stretches have variable thickness, the second border (in this case, outer border) is more difficult to find and fit. For this purpose, we identified two intensity minima: a first one that directly follows the first border, and a subsequent one (green arrows in panel 6). The strong intensity peak in between those minima is identified as the second border, and its position approximated by a Gaussian function. After WGA profiles from the whole data set have been analyzed, the mean calyx thickness was determined. All WGA profiles (7) as well as profiles of the analyzed proteins of interest (8) were averaged and scaled and aligned according to the mean calyx thickness. (**b**) For the analysis of AZ-specific protein distribution, two line profiles in AZ-negative (AZ(-)) and two profiles in AZ-positive (AZ(+)) presynaptic regions were drawn per calyx (using the algorithm described in **a**). (**c**) For colocalization analysis, line profiles were truncated to specifically cover only the presynaptic area and converted to area selections (1). Using a custom-written ImageJ plugin, all possible combinations of targets underwent Pearson’s colocalization analysis (2). All data were imported to Matlab to generate colocalization matrices (3), which were imported to Excel where they were averaged and finalized. (**a**-**c**) Scale bars correspond to 1 µm.

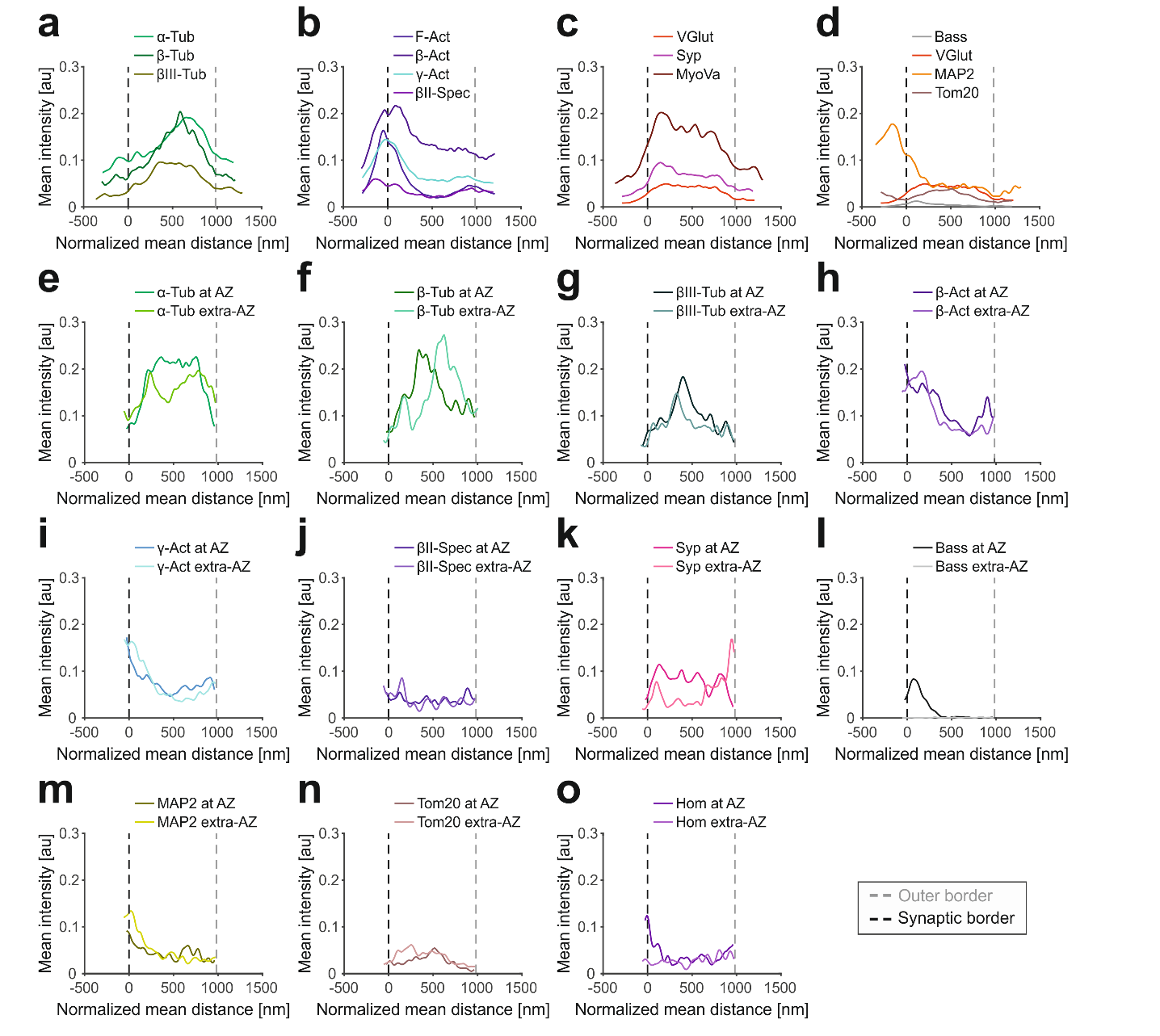

**Supplementary Figure 8 | Averaged line profiles of global and active zone-specific protein distributions.** (**a**) All analyzed tubulin (Tub) family members show a similar distribution. (**b**) Labels against polymerized actin (Act) or against α- and β-actin isoforms show similar profiles. Interestingly, βIII-spectrin distribution resembles that of actin. (**c**) Motor protein myosin Va is distributed throughout the presynapse in line with the distribution of its cargo reflected by synaptic vesicle markers VGlut and synaptophysin (Syp). (**d**) Different markers reflecting the overall geometry of the calyx of Held. (**e**-**o**) Distribution of proteins in active zone (AZ) positive versus AZ-negative calyx regions.

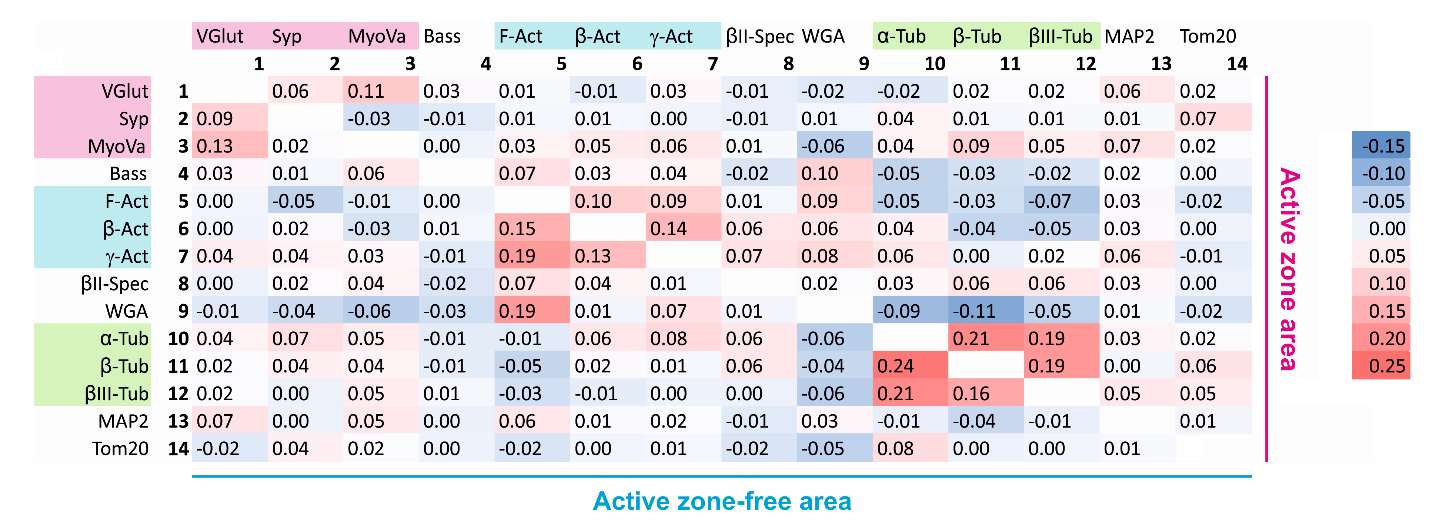

**Supplementary Figure 9 | Colocalization matrix.** Pearson’s r values for the colocalization matrix shown in **Figure 3m**.

**Supplementary Note 1**

*Extended information relating to control experiments assessing signal removal after each staining round (****Supplementary Fig. 3a****)*

We thoroughly assessed the performance of re-staining, bleaching, and elution with the following workflow, using Tom20 staining as a test case. For each control condition (**Supplementary Fig. 3a**), three *d*STORM imaging experiments (with three acquisitions per experiment and three ROIs per acquisition) were conducted, from which we quantified the fluorescence intensity by determining the number of single-molecule localizations. In between single imaging experiments, we eventually integrated an additional step, i.e. photobleaching the sample, elution of antibodies, or re-staining with a secondary antibody.

First, we determined the fluorescence intensity in three consecutive *d*STORM experiments performed with identical imaging conditions (first data set in **Supplementary Fig. 3a**). We quantified the fluorescence intensity per image by determining the number of localizations, and we normalized the numbers obtained for the second and third experiment to the first one. Under our experimental conditions, we find the expected decrease in fluorescence intensity caused by photobleaching of some of the fluorophore labels (see **Supplementary Fig. 3b** for graphical illustration).

Second, we performed an experiment with three consecutive *d*STORM acquisitions, but labeled again with the same fluorophore-labeled secondary antibody prior to the third imaging round (second data set). We find similar fluorescence intensities as in the first data set, despite re-labeling with the secondary antibody. This result tells us that the secondary antibody labeling reached saturation.

Third, we recorded three *d*STORM data sets, yet now photobleached the remaining fluorescence signal after the first acquisition by applying high irradiation intensities (see **Online Methods**). The second imaging round showed us that bleaching was efficient since the fluorescence intensity dropped to background level. Prior to the third experiment, we relabeled the sample again with the same fluorophore-labeled secondary antibody. We found a recovery of the fluorescence intensity, which we explain by photo-induced unbinding of the secondary antibody through high-intensity illumination of the sample during the photobleaching step. We find similar ratios of recovery (about 45%) as reported in the literature^6^.

Fourth, we explored the efficiency of elution (see **Online Methods**). After the first imaging round, we eluted the antibodies, and we observed a decrease in fluorescence intensity. A third imaging round shows a decreased intensity as observed in condition one and two.

Fifth, we explored the efficiency of both photobleaching and elution after the first imaging round. After the initial staining, we first exposed the sample to the elution buffer followed by a photobleaching step. We found the fluorescence intensity to drop to background level. Re-staining with the same secondary antibody recovers fluorescence signal (to ~7.5%), which we attribute to photo-unbinding of the secondary antibody^6^.

Sixth, we show that the combination of photobleaching and elution is highly efficient, if another primary antibody (other than Tom20; in this case Fibrillarin) competing for the same secondary antibody is introduced before the last acquisition round. This would represent the optimal workflow of a multiplex experiment.

In summary, for optimal performance of our approach, we recommend to design a re-labeling experiment such that (i) both elution and photobleaching steps are integrated in the workflow, and that (ii) the labeling sequence is designed such that the species of secondary antibodies changes every imaging round.

**Supplementary Note 2**

*Supplementary discussion of registration precision between different staining rounds for cells or tissues*

The serial re-staining approach used here requires a high-fidelity registration of images from each staining round to reliably derive conclusions on the positioning of each protein with regard to all other proteins investigated. We achieved this by imaging fiducial beads positioned between the sample and the coverslip for every staining round. Hence, assuming an invariant position of each bead, the registration procedure should work with similar precision for images acquired in any staining round. Protein positions determined from the first staining round should be equally comparable to those of the second or 10^th^ staining round. However, if the position of the fiducial beads varies to a small extent during the long duration of a typical multiplexed imaging experiment, deviations will occur, in particular for protein positions determined from imaging rounds far apart, but not much affecting neighboring staining rounds. This relationship is quite evident in **Figures 2b** and **3d.** Noticeably, the registration precision was overall better for super-resolution images acquired in tissue (**Fig. 3d**) compared to cells (**Fig. 2b**). We attribute this to a differential stability of the fiducials in these two imaging situations as detailed in the following. In contrast to the flat 400 nm sections, cultured U2OS cells have a substantial thickness of several micrometers. Due to their thickness and other mechanical properties, cells may not homogeneously adhere to the glass surface, resulting in a slight partial detachment with each additional staining round. This could in turn loosen fiducial beads stuck underneath the cells. Slight changes in cell volume with increasing staining and elution rounds may cause a similar effect. Therefore, to re-establish a focal plane in late imaging rounds comparable to that from the initial round, the focus would have to be placed more and more deeply into the sample, resulting in less focused fiducial beads, and thus, less precise fitting and a higher registration error. In the much more homogeneous tissue sections, these effects will impact registration precision much less, consistent with the experimental data (**Figs. 2b** and **3d**). Hence, we have focused our quantitative analyses on neuronal tissue samples to minimize these complications. Moreover, for the analysis of protein relationships we performed five multiplex experiments with target proteins imaged in a different order. Thereby, we balanced out possible effects arising from misalignment in late compared to early imaging rounds. Future work may include the use of fiducials covalently fixed to the cover slip using nano-engineering procedures.
